# MYT1L is required for suppressing earlier neuronal development programs in the adult mouse brain

**DOI:** 10.1101/2022.10.17.512591

**Authors:** Jiayang Chen, Nicole Fuhler, Kevin Noguchi, Joseph D. Dougherty

**Author notes:** Corresponding author, Dr. Joseph D. Dougherty, Washington University School of Medicine, Department of Psychiatry, Department of Genetics, Campus Box 8232, 660 South Euclid Avenue, St. Louis, MO 63110-1093.

## Abstract

*In vitro* studies indicate the neurodevelopmental disorder gene Myelin Transcription Factor 1 Like (MYT1L) suppresses non-neuronal lineage genes during fibroblast-to-neuron direct differentiation. However, MYT1L’s molecular and cellular functions during differentiation in the mammalian brain have not been fully characterized. Here, we found that MYT1L loss leads to up-regulated deep layer (DL) but down-regulated upper layer (UL) neuron gene expression, corresponding to an increased ratio of DL/UL neurons in mouse cortex. To define potential mechanisms, we conducted Cleavage Under Targets & Release Using Nuclease (CUT&RUN) to map MYT1L binding targets in mouse developing cortex and adult prefrontal cortex (PFC), and to map epigenetic changes due to MYT1L mutation. We found MYT1L mainly binds to open chromatin, but with different transcription factor co-occupancies between promoters and enhancers. Likewise, multi-omic dataset integration revealed that, at promoters, MYT1L loss does not change chromatin accessibility but does increase H3K4me3 and H3K27ac, activating both a subset of earlier neuronal development genes as well as *Bcl11b*, a key regulator for DL neuron development. Meanwhile, we discovered that MYT1L normally represses the activity of neurogenic enhancers associated with neuronal migration and neuronal projection development by closing chromatin structures and promoting removal of active histone marks. Further, we show MYT1L interacts with SIN3B and HDAC2 *in vivo*, providing potential mechanisms underlying any repressive effects on histone acetylation and gene expression. Overall, our findings provide a comprehensive map of MYT1L binding *in vivo* and mechanistic insights to how MYT1L facilitates neuronal maturation.

## Introduction

Neuronal development is a continuous process starting early during embryogenesis and lasting well into the postnatal ages (Kroon et al. 2019; Stiles and Jernigan 2010). Originating from asymmetric divisions of neural progenitors in the ventricular zone (VZ), neurons undergo earlier neuronal development programs, including migration and projection development, followed by later maturation processes, including synaptic development and pruning, to gain their locational and functional identities in different layers of the cortex (Campbell 2005; Götz and Huttner 2005; Luo and O’Leary 2005). These steps are finely tuned by a sophisticated network of cis-regulatory elements (e.g., promoters and enhancers), trans-regulatory factors (e.g., transcriptional factors(TFs) and histone modifying complexes), as well as epigenetic regulation (e.g., changes in chromatin accessibility and histone modifications) (Dixit et al. 2014; Lomvardas and Maniatis 2016; Nitarska et al. 2016; Olson et al. 2001; Trevino et al. 2020; Yousefi et al. 2021). Thus, proper activation and repression of those early neuronal development programs in a timely manner is crucial for both generation of neurons and their later maturation and fate specification (Yuan et al. 2022). TFs, especially proneuronal basic helix-loop-helix (bHLH) TFs, are key components that orchestrate these programs (Dixit et al. 2014; Olson et al. 2001; Tutukova et al. 2021). Several master bHLHs, such as NEUROG2 and NEUROD1, are expressed at different developmental stages and are responsible for facilitating neuronal fate progression by modulating promoters, enhancers, and thus their corresponding genes’ expression (Dixit et al. 2014; Noack et al. 2022; Pataskar et al. 2016). Although multi-omics dataset integration has helped identify many of these cis- and trans-regulatory elements involved in neuronal development, how their activities are precisely controlled to produce and mature neuronal cell types remains poorly understood.

In addition to bHLHs, a variety of additional TFs, including TBR1 and BCL11B, are also developmentally expressed in a cell-type specific manner, and play indispensable roles in neuronal development (Arlotta et al. 2008; Bedogni et al. 2010). Myelin Transcription Factor 1 Like (MYT1L), a pro-neuronal TF expressed primarily in postmitotic neurons, appears to be one such key factor participating in neuronal fate specification (Heavner et al. 2020; Mall et al. 2017) and maturation (Chen et al. 2021; Mall et al. 2017). For example, *in vitro* studies by shRNA knockdown suggested that MYT1L loss increases the ratio of deep layer (DL) to upper layer (UL) cortical neurons (Heavner et al. 2020). Furthermore, overexpressing MYT1L, along with ASCL1 and BRN2, can directly reprogram mouse embryonic fibroblasts (MEFs) into functional neurons, demonstrating its potent role in promoting neuronal differentiation (Vierbuchen et al. 2010). Utilizing the same reprogramming system, Mall et. al mapped MYT1L targets by Chromatin Immunoprecipitation followed by Deep Sequencing (ChIP-seq) and measured target gene expression by RNA-seq (Mall et al. 2017). Interestingly, in this system MYT1L mainly acts as a transcriptional repressor that silences non-neuronal gene expression to facilitate neuronal differentiation (Mall et al. 2017). Meanwhile, other *in vitro* studies revealed that MYT1L can function as both a transcriptional activator and repressor, probably through distinct protein domains. For example, truncation experiments have shown the N-terminal activates gene expression in reporter assays, while a central zinc-finger containing domain suppresses expression (Manukyan et al. 2018). However, in the transdifferentiation system, the N-terminus together with central zinc-finger domains recruit co-factors including SIN3B and are sufficient for neuronal reprogramming (Mall et al. 2017). SIN3B is thought to then recruit histone deacetylases (HDACs) (Bainor et al. 2018), although direct interactions between MYT1L, SIN3B and HDACs have not been shown *in vivo*, nor has MYT1L’s impact on histone modifications been examined. Overall, despite widespread usage of MYT1L in neuronal transdifferentiation, the molecular mechanisms underlying MYT1L’s pro-neuronal activities during normal brain development remain incompletely defined. Additionally, it is unclear whether MYT1L influences epigenetic landscapes to regulate gene expression.

Recently MYT1L has also been implicated in human neurodevelopmental disorder (NDD), with the spectrum of phenotypes caused by MYT1L loss of function (LoF) mutations now recognized as MYT1L Syndrome (Blanchet et al. 2017; Coursimault et al. 2021). To understand MYT1L’s functions *in vivo* and how MYT1L mutations lead to human disease pathology, several *in vivo* models have been established (Blanchet et al. 2017; Chen et al. 2021; Mall et al. 2017; Wöhr et al. 2022). Earlier studies showed, MYT1L homologs’ (myt1la and myt1lb) knockdown in zebrafish by antisense morpholinos reduces oxytocin and arginine vasopressin mRNA abundance in hypothalamus, suggesting MYT1L is important for neuroendocrine system development, and/or neuronal maturation, as peptide expression develops relatively late postnatally (Almazan et al. 1989; Blanchet et al. 2017). Furthermore, MYT1L shRNA knockdown by *in utero* electroporation impairs neuronal migration in the developing mouse cortex, echoing its roles in facilitating neuronal development (Mall et al. 2017). As MYT1L Syndrome patients harbor *de novo* heterozygous mutations of MYT1L, more recently a MYT1L germline knockout mouse line was generated to mimic human patient genetics (Chen et al. 2021). This study showed that MYT1L heterozygous knockout (Het) mice recapitulate many phenotypes reminiscent of the human syndrome, including obesity, hyperactivity, and social deficits. Key phenotypes were replicated in two additional MYT1L haploinsufficiency mouse models (Wöhr et al. 2022; Kim et al. 2022). In the initial MYT1L germline knockout mouse line, it was also demonstrated that MYT1L loss results in insufficient cell proliferation in embryonic mouse cortex (Chen et al. 2021). Further investigation leveraging existing ChIP-seq data (albeit from *in vitro* binding experiments (Mall et al. 2017)), along with *in vivo* ATAC-seq, and RNA-seq datasets revealed MYT1L has a role in activating cell proliferation programs but suppressing early neural differentiation programs. In contrast with predictions of the transdifferentiation system, no obvious activation of non-neuronal lineage genes was found in Het mice (Chen et al. 2021). Yet, one limitation of these analyses was they were based on ChIP-seq from an orthogonal system. Since ectopic overexpression of MYT1L during transdifferentiation might distort normal MYT1L binding activity, high quality binding profiles of MYT1L *in vivo* are needed to better understand its functions in physiological conditions. In addition, even though there is sustained expression of MYT1L in the adult brain (Chen et al. 2021), little was known about its functions and binding in later stages of neuronal development, and long-term consequences of MYT1L loss have not been assessed.

Therefore, we adopted CUT&RUN technology to define MYT1L binding targets in the cortex and investigated its molecular functions through epigenetic profiling and further neuroanatomical studies *in vivo*. First, we found that MYT1L loss alters DL and UL neuron numbers in adult mouse cortices, consistent with the hypothesis emerging from the previous *in vitro* study. We then optimized CUT&RUN profiling using knockout (KO) controls in embryos, and then mapped MYT1L binding in adult mouse prefrontal cortex (PFC). Simultaneous assessment of epigenetic changes revealed that MYT1L regulates promoter and enhancer chromatin accessibility and active histone modifications in the PFC, potentially via interactions with SIN3B and HDAC2. These epigenetic modifications provide an appropriate level of suppression of early neuronal development programs in the adult brain, including those for earlier processes like neuronal migration and neuronal projection development, a suppression that is lost in the Hets. In addition, we found MYT1L loss directly activates *Bcl11b* expression, which could explain the increased DL neuron number in the Het cortex. In sum, this study unravels MYT1L’s role in repressing earlier neuronal development programs during the later stages of neuronal maturation *in vivo* and provides insight into long-term consequences of MYT1L loss during normal brain development.

## Results

### MYT1L loss alters the ratio of deep/upper layer neurons in mouse cortex

Previous *in vitro* study has shown MYT1L knockdown increased the ratio of DL/UL neurons (Heavner et al. 2020). Thus, we first investigated whether MYT1L constitutive Het knockouts, mimicking the gene dose of patients, can also result in similar phenotypes in mice. First, we re-evaluated the RNA-seq dataset from MYT1L mutant (Het and KO) embryonic day 14 cortex (E14 CTX) and MYT1L Het PFC (Chen et al. 2021). With Gene Set Enrichment Analysis (GSEA), we found there is a significantly increased expression of DL neuron signature genes in mutant E14 CTX compared with wild-type (WT) (**Fig 1A**), while expression of the UL neuron signature genes is unchanged (**Fig 1B**); however as UL genes are not yet highly expressed by E14, this finding may not be conclusive. Therefore, we also examined adult PFC when both types are present. In the adult Het PFC, the expression of DL genes is even more significantly up-regulated (**Fig 1C**), and UL genes demonstrate significant down-regulation (**Fig 1D**), suggesting that the impact of MYT1L loss on DL and UL gene expression is not a transient effect. As DL neurons are in an earlier neuronal development trajectory than UL upper layer neurons, such an up-regulation of DL neuron genes upon MYT1L loss is consistent with the hypothesis that normal MYT1L levels are needed to facilitate neuronal maturation.

**Figure 1:**
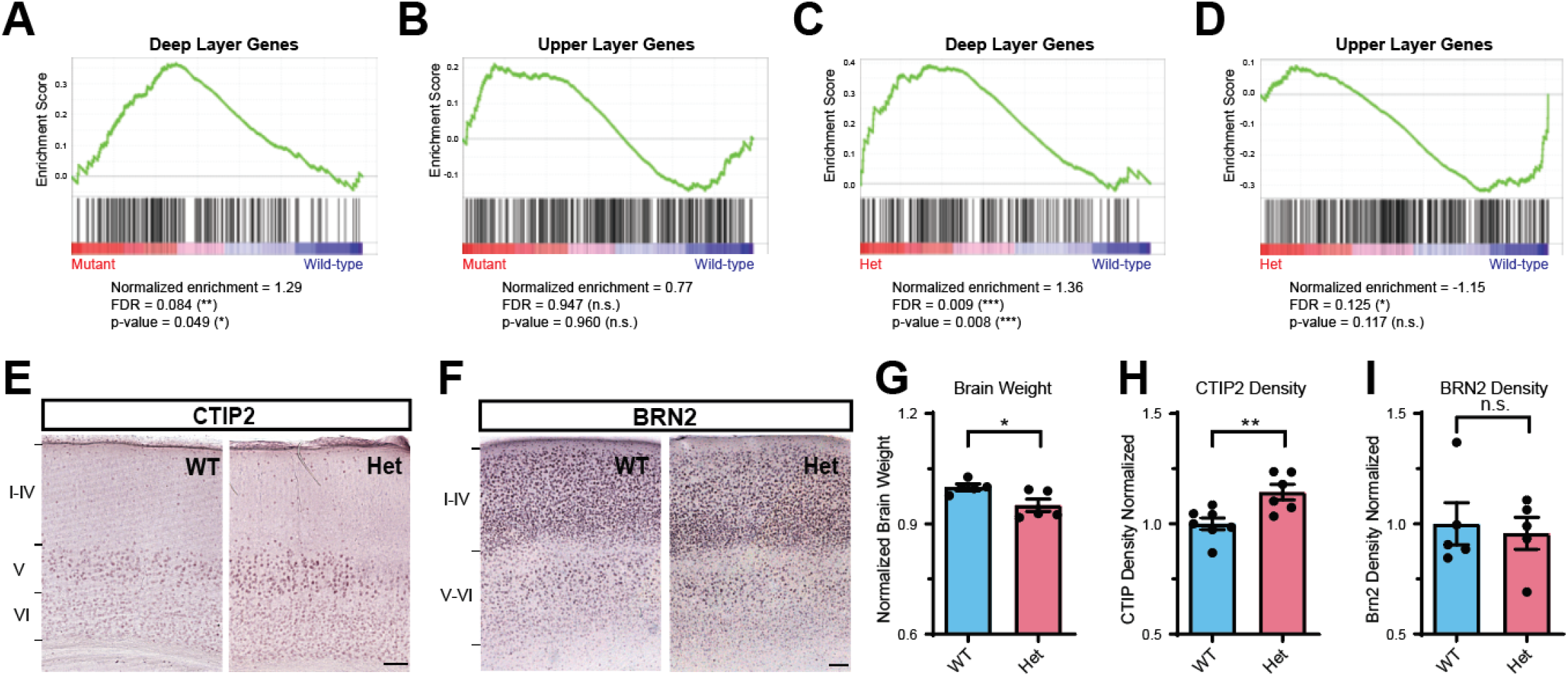
MYT1L controls cortical neuron layer specification. (**A**) GSEA showed an up-regulation of DL genes in MYT1L mutant E14 CTX. (**B**) UL genes showed no significant change in MYT1L mutant E14 CTX. (**C**) GSEA showed an up-regulation of DL genes in MYT1L Het P60 PFC. (**D**) UL genes showed subtle but significant down-regulation in MYT1L Het P60 PFC. (**E**) Representative images of CTIP2 staining on the P60 mouse cortex. (**F**) Representative images of BRN2 staining on the P60 mouse cortex. (**G**) MYT1L Het mice have reduced brain weights compared to WTs. (**H**) MYT1L Het mice have increased CTIP2+ neuron density in cortex. (I) BRN2+ neuron density remains the same between Hets and WTs. *p<0.05, **p<0.01. Data were represented as Mean±SEM.

One possible explanation for the adult RNA-Seq pattern is that Hets have more DL neurons than wild-type controls. Thus, to test this hypothesis and examine MYT1L’s role in regulating neuronal localization *in vivo*, we stained the postnatal day 60 (P60) Het cortex (**Fig 1E,F**) with UL and DL markers. As expected, we replicated the finding that Het mice have reduced brain weights compared with WT littermates (**Fig 1G**)(Chen et al. 2021). Consistent with our hypothesis, we found Het cortices have increased DL neuron density (labeled by CTIP2 protein, encoded by the *Bcl11b* gene) compared with WT littermates (**Fig 1E, H**). On the other hand, UL neurons (labeled by BRN2, encoded by the *Pou3f2 gene*) did not show altered density in Het cortices (**Fig 1F, I**). In the RNA-seq, MYT1L loss did incur much smaller effects on upper layer neuron genes (**Fig 1B,D**), which might not be distinguishable by immunochemistry experiments. In sum, both RNA-seq and immunochemistry validation experiments showed MYT1L loss also altered the ratio of DL/UL neurons in the mouse cortex.

### CUT&RUN identifies MYT1L binding targets in the mouse embryonic cortex

To understand how MYT1L regulates the DL/UL neuron ratio, as well as the altered transcriptional profile we previously observed (Chen et al. 2021), we next investigated MYT1L’s molecular functions *in vivo*. MYT1L has peak protein expression between E14 and P1 in the mouse brain (Chen et al. 2021). In order to map MYT1L targets *in vivo*, we optimized CUT&RUN on E14 mouse cortex (**Fig 2A**) (Skene and Henikoff 2017). First, leveraging the MYT1L germline knockout (S710fsX) mouse line (Chen et al. 2021), we validated CUT&RUN and antibody specificity on MYT1L KO samples. These S710fsX mice do not produce any MYT1L protein and thus can serve as a gold-standard control.

**Figure 2:**
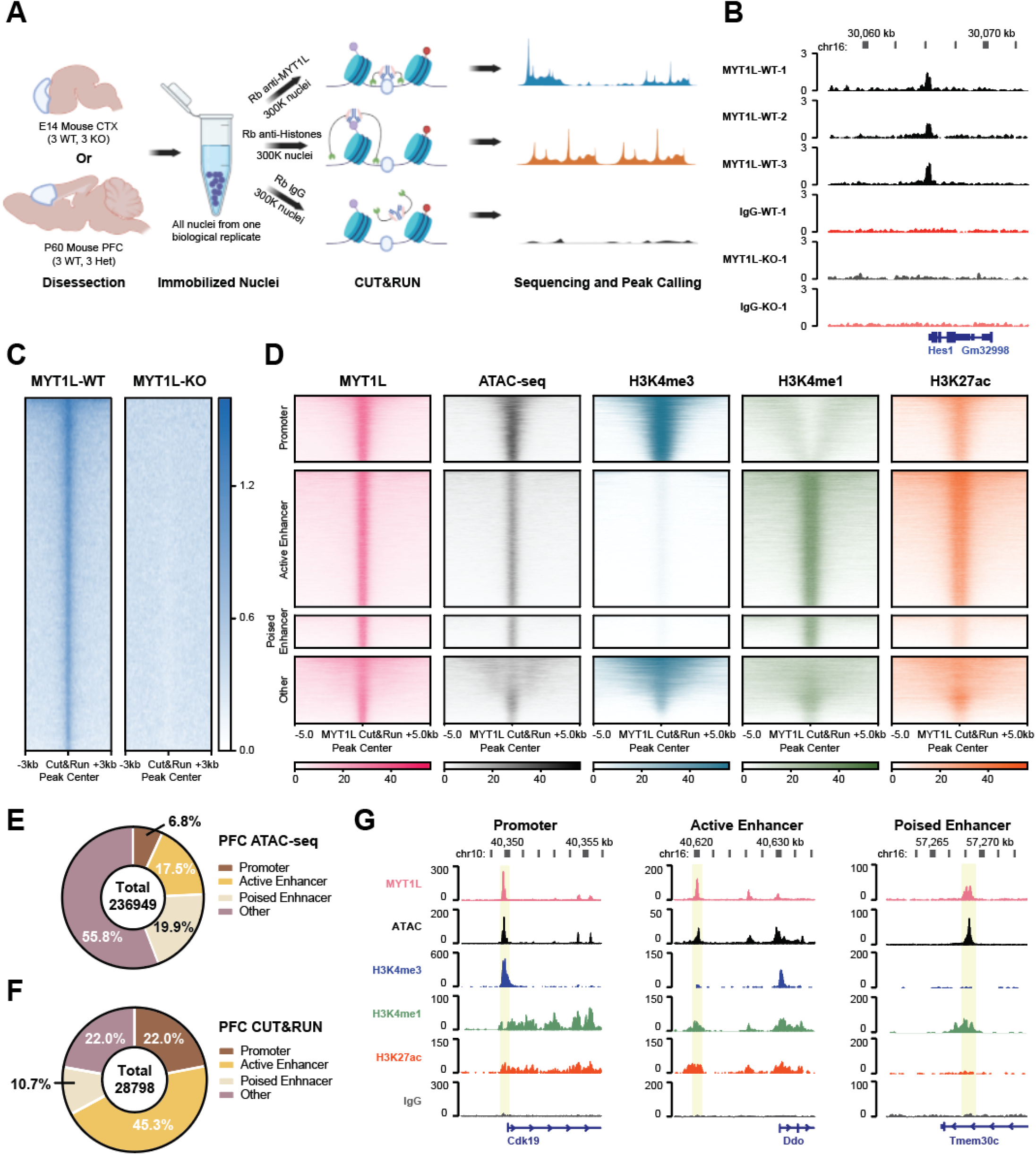
CUT&RUN identifies MYT1L specific binding targets in the E14 mouse cortex. (**A**) Workflow of CUT&RUN experiments on E14 CTX and adult prefrontal Cortex (PFC). (**B**) Representative IGV tracks showing a reproducible MYT1L peak at the Hes1 promoter region in all 3 WT E14 biological replicates but not in IgG and KO samples. (**C**) Heatmaps of 20,305 MYT1L peaks called from the merged bam file. MYT1L CUT&RUN signals were diminished in the KO. (**D**) Heatmaps of CUT&RUN signals of MYT1L, IgG, and histones at MYT1L bound regions in PFC. (**E**) Annotations of PFC ATAC-seq peaks show the genome-wide distribution of promoters, active enhancers, and poised enhancers in open chromatin regions. (**F**) Annotations of MYT1L targets in PFC show MYT1L mainly binds to active enhancers. (**G**) Representative genome tracks of MYT1L bound promoter (left), active enhancer (middle), and poised enhancer (right).

Next, we compared two peak calling methods, a stringent one where we called peaks from each biological replicate independently and intersected peak profiles from all three, and a more sensitive approach where we called peaks from a single merged alignment file of all three samples. In the intersected peaks, we identified 560 MYT1L peaks in wild-type (WT) E14 mouse cortex, and no peaks in all KO samples (**Fig 2B, Table S1**), indicating specificity. With the increased sensitivity approach, we identified 20305 unique peaks in WT compared with KO (**Fig 2C, Table S1**). Although MYT1L binding activity was still absent in KOs (**Fig 2C**), this method led to a higher percentage of lower-enrichment peaks (49.1% of targets with enrichment score < 3, 9969 out of 20305, **Fig S1A**). In addition, in *de novo* motif finding, the known MYT1L core binding motif AAGTT (Jiang et al. 1996; Mall et al. 2017) was significantly enriched in all 560 intersected peaks (100% of targets, *p* = 1e-11, **Fig S1B,C**) but not in all peaks called from merged alignment file (**Fig S1E**), suggesting peak calling from merged files might be recovering either more low-affinity bindings or experimental noise. Thus, we provide both high and low stringency tables for future use. Regardless, since MYT1L binding profiles have not been well characterized after development, we next conducted CUT&RUN the adult mouse brain. Also, E14 brain has relatively few MYT1L-expressing cells (neurons), thus adult brain with its higher neuron proportion may map MYT1L binding in a more efficient way.

### Using adult mouse prefrontal cortex improves CUT&RUN sensitivity on MYT1L profiling

To better compare with existing multi-omics datasets and human phenotypes, we chose adult PFC as the target region for MYT1L CUT&RUN. This brain region is associated with attention deficit and hyperactivity disorder (ADHD) (Yasumura et al. 2019). ADHD is observed in MYT1L Syndrome human patients, and hyperactivity is found in the mouse models. We again compared two peak calling methods. First, we identified 28,798 reproducible MYT1L bound peaks by intersecting peak calls from 3 biological replicates of WT mouse PFC (**Table S1, Fig S1C**), and the MYT1L core binding motif AAGTT is significantly enriched via de novo motif finding (76.37% of targets, *p* = 1e-3125) (**Fig S1C, D**). Next, we did peak calling from the merged alignment file. This resulted in 115,143 peaks with slightly improved peak enrichment (55.7 % > 3 vs. 50.9% > 3 in E14) compared with the E14 dataset (**Fig S1A**, **Table S1**). Motif analysis also showed a significantly enriched AAGTT motif (33.38% of targets, *p* = 1e-713) (**Fig S1E,F**). Overall, many more peaks were identified from PFC CUT&RUN experiments, yet the majority of the E14 peaks were also recovered in PFC CUT&RUN (**Fig S1C,E**), suggesting CUT&RUN on PFC is more efficient than E14. In addition, we compared our MYT1L CUT&RUN targets with MYT1L ChIP-seq data from both E14 brain and fibroblasts overexpressing MYT1L, BRN2, and ASCL1 (Mall et al. 2017). Surprisingly, we did not see substantial overlap between MYT1L CUT&RUN and ChIP-seq peaks (**Fig S2A-D**). Notably, MYT1L CUT&RUN peaks overlap better with MYT1L brain ChIP-seq than with MYT1L fibroblast ChIP-seq (**Fig S2A-D**), suggesting MYT1L binding is very context dependent with different binding in different cell types, and thus MYT1L likely does not serve as a pioneer factor that opens the chromatin of its targets in any cellular context. Since intersected peaks from PFC have the best MYT1L motif enrichment, we focused on the 28,798 MYT1L high-stringency binding targets identified from PFC CUT&RUN for the downstream analysis to understand MYT1L’s functions in the adult brain and the long-term consequences of MYT1L loss.

### MYT1L co-occupies its binding sites with different transcription factors at promoter and enhancer regions

Previous ChIP-seq experiments have shown that MYT1L mainly binds to the promoter regions when overexpressed during reprogramming of MEFs (Mall et al. 2017). To test if it is also true *in vivo*, we annotated MYT1L targets from adult mouse PFC. Analysis of MYT1L of colocalization at these regions showed that MYT1L tends to bind open chromatin regions (95.3%, 27450/28798) with enhancers being the most common category, when compared to the categories of all open chromatin regions (**Fig 2E,F, Table S2**). Meanwhile, we utilized nuclei from the same animals to perform CUT&RUN on several histone modifications, including H3K4me3, H3K4me1, and H3K27ac (**Fig 2A**). Leveraging these histone modification profiles, we further categorized enhancers (**Table S3**) into poised (H3K4me1+/H327ac-) and active (H3K4me1+/H3K27ac+) as previously described (Creyghton et al. 2010). To understand sequence preferences of MYT1L at promoters, poised enhancers, and active enhancers, we performed motif analysis using monaLisa to compare *de novo* binding motifs and predicted TFs co-occupancy between the three. Surprisingly, we found motifs for TFs that behave as transcriptional activators’ motifs (e.g., SP1 and ELK1) to be enriched in MYT1L bound promoters, while neurogenic TFs (e.g., MEF2A and NEUROD1) and activity dependent TF (e.g., JUNB) motifs were specifically enriched in MYT1L bound enhancers (**Fig 3A**), even when controlling for the differential baseline frequency of these TF motifs at promoters, poised enhancers and active enhancers respectively. Furthermore, this is not driven by the MYT1L core binding motif AAGTT since both MYT1L bound promoters and enhancers are significantly enriched for AAGTT (**Fig 3B**). In order to investigate if these motifs abundances reflected the actual TF co-occupancy, we compared MYT1L CUT&RUN targets with published TF ChIP-seq data. As expected, compared to MYT1L unbound regions (MYT1L-), ChIP-seq peaks of candidate TFs, including SP1, ELK1, MEF2A, JUNB, NEUROD1, and NEUROD2, are significantly enriched in MYT1L bound genomic regions (MYT1L+) (**Fig S3A-G**), suggesting these TFs are frequently bound at MYT1L bound peaks. Next, we compared TF ChIP-seq enrichment between the MYT1L+ promoters and the MYT1L+ enhancers. Echoing the motif analysis, MYT1L has significantly higher co-occupancy with SP1 and ELK1 in promoter regions than in enhancer regions (**Fig 3C,D, S3G**). Likewise, MEF2A and JUNB prefer binding MYT1L enhancer targets over promoter targets (**Fig 3E,F, S4G**). However, in contrast to the motif analysis, a much higher percentage of bHLH TFs NEUROD1 and NEUROD2 ChIP-seq peaks are at MYT1L-bound promoter targets than at MYT1L bound enhancers (**Fig 3G,H, S3G**), suggesting relative motif enrichment will not always predict the corresponding TF binding, or that a different protein might be binding this sequence at enhancers. Meanwhile, MYT1L does not appear to block the binding of NEUROD1 and NEUROD2 at enhancers since no obvious depletion of TF binding in MYT1L+ enhancers was observed (**Fig S3A-F**). Overall, such differential TF co-occupancy suggests MYT1L might play different roles at promoters and enhancers by cooperating with different co-factors.

**Figure 3:**
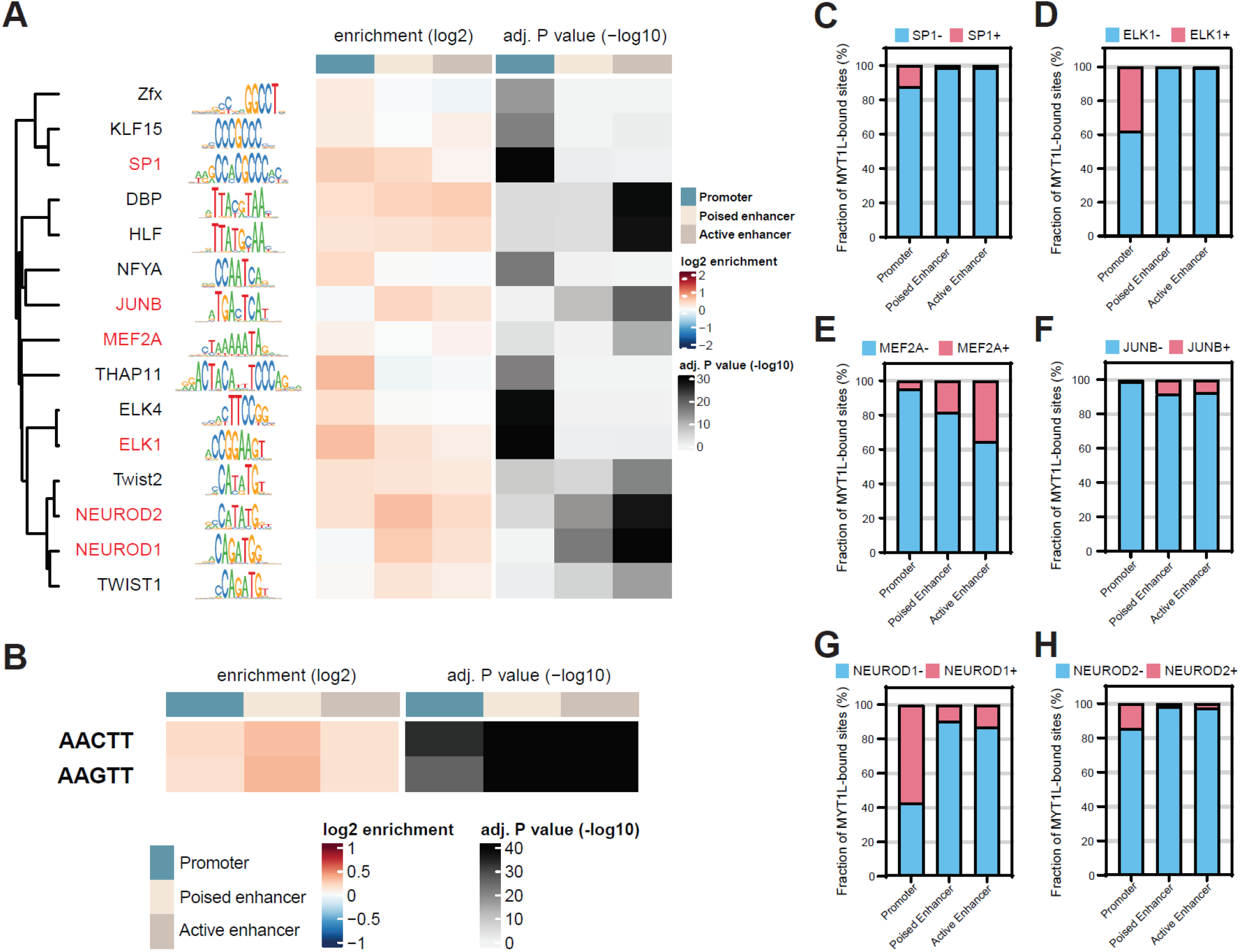
MYT1L co-occupies with different sets of TFs at promoter and enhancer regions. (**A**) monaLisa motif analysis revealed that MYT1L co-occupies with transcriptional activators such as ELK1 at promoter regions, while it co-occupies with neurogenic TFs such as MEF2A at enhancer regions. (**B**) Both MYT1L bound promoters and enhancers are significantly enriched for MYT1L core binding motif, AAGTT. (**C**) Overlapping between MYT1L CUT&RUN targets and TFs ChIP-seq peaks showed that more MYT1L promoter targets were also bound by transcriptional activators like SP1 and (**D**) ELK1 than enhancer targets, while more enhancer targets were bound by (**E**) the neurogenic TF MEF2A and (**F**) activity dependent gene JUNB. (**G**) NEUROD1 and (**H**) NEUROD2 have stronger presence at MYT1L promoter targets than enhancer targets.

### MYT1L directly binds to promoters of genes involved in early neuronal development

To understand molecular functions of MYT1L binding, we first focused on MYT1L bound promoters since they can be directly associated with nearby genes, thus allowing functional inferences of downstream consequences. Chromatin accessibility is closely correlated with gene expression, where open chromatin indicates active gene expression while closed chromatin indicates gene repression. Therefore, we compared MYT1L binding data with ATAC-seq (Assay of Transposase Accessible Chromatin sequencing) on P60 mouse PFC (Het versus WT) (Chen et al. 2021) to see whether MYT1L binding changes chromatin structures at promoters. We found that MYT1L haploinsufficiency indeed reduces overall MYT1L binding activities, but does not affect chromatin accessibility at promoter targets (**Fig 4A,B, S4A**). Histone modifications are also closely correlated with gene expression, where H3K4me3 and H3K27ac are thought to be marks for active gene expression (Black et al. 2012; Heintzman et al. 2009). Thus, we investigated how histone landscape changes at promoter regions upon MYT1L loss. We found there were more H3K4me3 and H3K27ac modifications in Het PFC at MYT1L promoter targets compared with WT (**Fig 4C,D, S4B,C**), indicating MYT1L’s role is normally to suppress gene expression by facilitating the removal of these marks. Previous *in vitro* studies have shown MYT1L can bind to SIN3B, a transcriptional repressor that recruits histone deacetylase (HDAC1/2) and demethylase (KDM5A) (Bainor et al. 2018; Naruse et al. 1999; Romm et al. 2005, 1; Hayakawa et al. 2007; Nishibuchi et al. 2014). We found MYT1L interacts with SIN3B as well as HDAC2 in the mouse cortex as well from co-immunoprecipitation (Co-IP) experiments, while no direct interaction between MYT1L and HDAC1 was observed (**Fig. 4E**). These results suggest, although MYT1L has minimal effects on chromatin accessibility at its bound promoters, it can facilitate the removal of active histone marks at promoters potentially via interacting with the SIN3B repressor complex.

**Figure 4:**
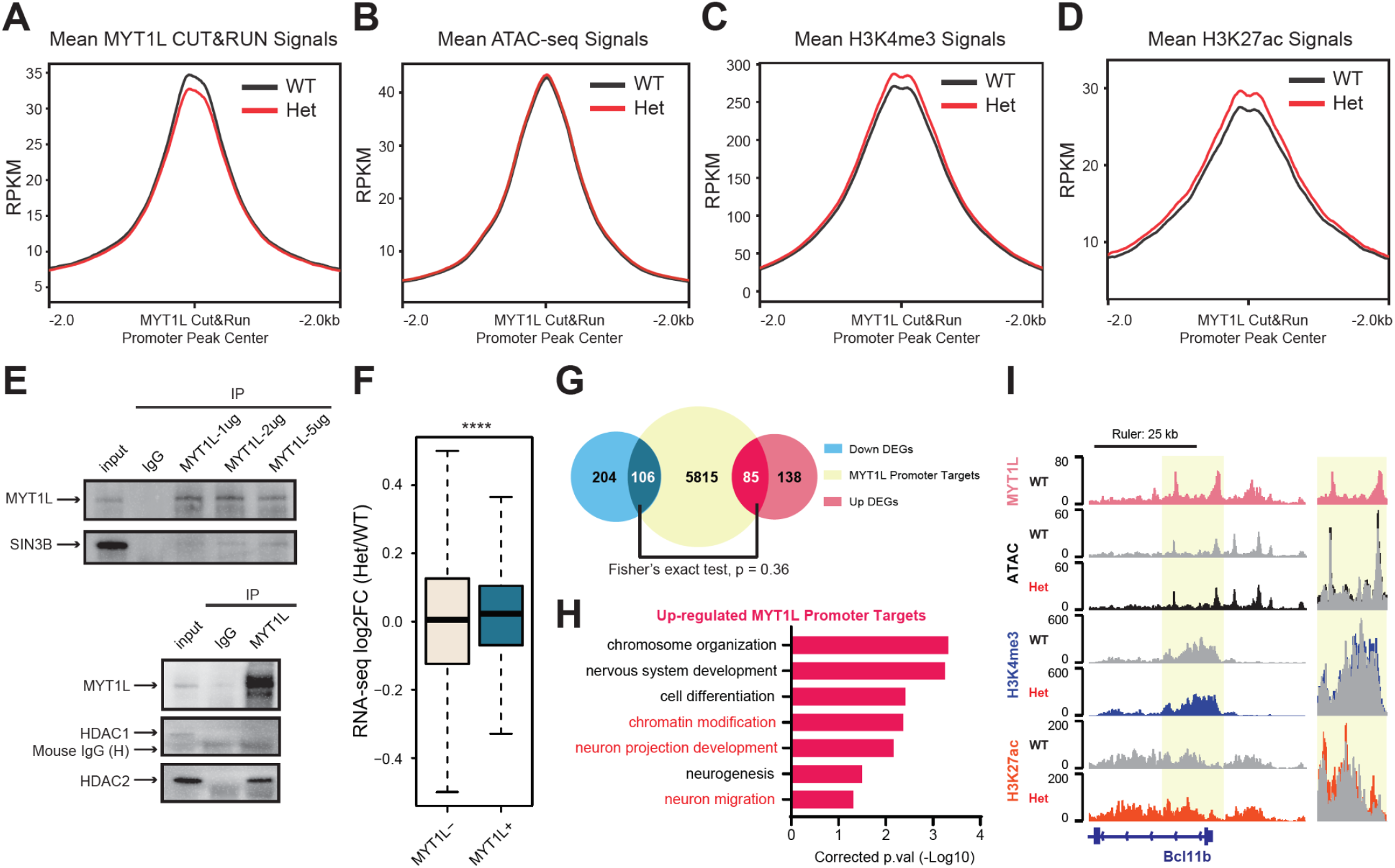
MYT1L directly binds to and controls promoters that are associated with early neuronal development genes. (**A**) Mean MYT1L CUT&RUN signals showed decreased MYT1L binding at promoters in Het PFC. (**B**) No chromatin accessibility change was observed at MYT1L bound promoters. (**C**) Mean H3K4me3 CUT&RUN signals showed increased H3K4me3 at MYT1L bound promoters in Het PFC. (**D**) Mean H3K27ac CUT&RUN signals showed increased H3K27ac at MYT1L bound promoters in Het PFC. (**E**) MYT1L co-immunoprecipitated with SIN3B and HDAC2 but not with HDAC1/3 in WT mouse cortex. (**F**) MYT1L loss increases its promoter targets’ expression in PFC. (**G**) Venn Diagram showing the overlaps among dDEGs, MYT1L Promoter targets, and uDEGs, and no biased overlap was observed (p = 0.36). (**H**) GO analysis on 85 uDEGs whose promoters are bound by MYT1L. (**I**) Representative genome browser track shows MYT1L-bound Bcl11b promoter has higher H3K4me3 and H3K27ac levels in Het PFC than WT. *p<0.05, **p<0.01, ***p<0.001, ****p<0.0001.

We next sought to determine the impact of these epigenetic changes at promoters on gene expression. By looking at adult Het PFC RNA-seq fold changes of genes whose promoters are bound by MYT1L, we saw a subtle but significant upregulated expression of those MYT1L targets in Het (**Fig 4F**), echoing MYT1L’s role as a transcriptional repressor in reprogramming neurons (Mall et al. 2017). Then, we specifically looked at differentially expressed genes (DEGs). Among 310 down-regulated genes (dDEGs) and 223 up-regulated genes (uDEGs) in MYT1L Het PFC, we saw no biased distribution of MYT1L promoter targets, with 106 of the dDEGs and 85 of the uDEGs’ promoters bound by MYT1L (**Fig 4G**). Gene ontology (GO) analysis on these overlapped genes revealed that MYT1L promoter targets up-regulated in the MYT1L Het are significantly enriched in chromatin modification (e.g., *Hdac4*, **Fig S4D**) and neuron projection development (e.g., *Lingo1* and *Cit*) pathways (**Fig 4H**). Notably, several key regulators of neuronal migration (e.g., *Ctnnd2*, **Fig S4E**) and deep cortical layer identity are directly bound by MYT1L and up-regulated in Het PFC, including *Bcl11b*, a master regulator of DL fate (Arlotta et al. 2005) (**Fig 4I**). However, mature neuron functional pathways, like synaptic transmission and ion transport, that were down-regulated in Het PFC are not enriched in MYT1L promoter targets (**Table S4**), indicating their dysregulation is likely an indirect effect of MYT1L loss. Together, these findings suggest that MYT1L directly suppresses neuronal development programs for chromatin, migration, and neurite extension by altering corresponding promoters’ epigenetics in the adult mouse cortex. In addition, MYT1L loss increases H3K4me3 and H3K27ac at promoters of DL marker genes, including *Bcl11b*, resulting in an up-regulated gene expression and increased DL neuron numbers.

### Loss of MYT1L activates enhancers associated with early neuronal development

Previous studies have shown enhancers are crucial for controlling neurodevelopmental programs as well as neuronal functions (Lu et al. 2020; Malik et al. 2014). Given enhancers have spatiotemporal activities throughout development (Carullo and Day 2019), and most of MYT1L targets identified in PFC are enhancers, we investigated how MYT1L binding influences enhancer epigenetics. After defining enhancers into active and poised stages by H3K27ac enrichment (**Fig 3A**), we found MYT1L preferentially binds to activated enhancers compared to poised enhancers in the adult mouse PFC (**Fig 3C**), and that the Het PFC has reduced MYT1L binding at both active and poised enhancers (**Fig S5A**). Next, we explored how MYT1L bound enhancers are developmentally regulated. By integrating the histone CUT&RUN data from E14 CTX, we defined active and poised enhancers in E14 CTX and compared those with adult PFC enhancers. Out of 13,050 MYT1L bound active enhancers in adult PFC, 80% (10,443/13,050) are adult-specific active enhancers and only 20% are shared at both developmental time points (**Fig 5A,B**). Furthermore, 24.8% (2,593/10,443) of those adult-specific active enhancer targets were poised in E14 CTX (**Table S2**), suggesting MYT1L might also guide the activation of a small subset of poised enhancers during development. On the other hand, out of 3077 MYT1L targets annotated as E14 CTX active enhancers, only 21.3% (656/3077) of them are E14 CTX-specific active enhancers (**Table S2**). Apparently, MYT1L occupancy at active enhancers is more prevalent in the adult stage, consistent with its major expression pattern in post-mitotic neurons (Chen et al. 2021).

**Figure 5:**
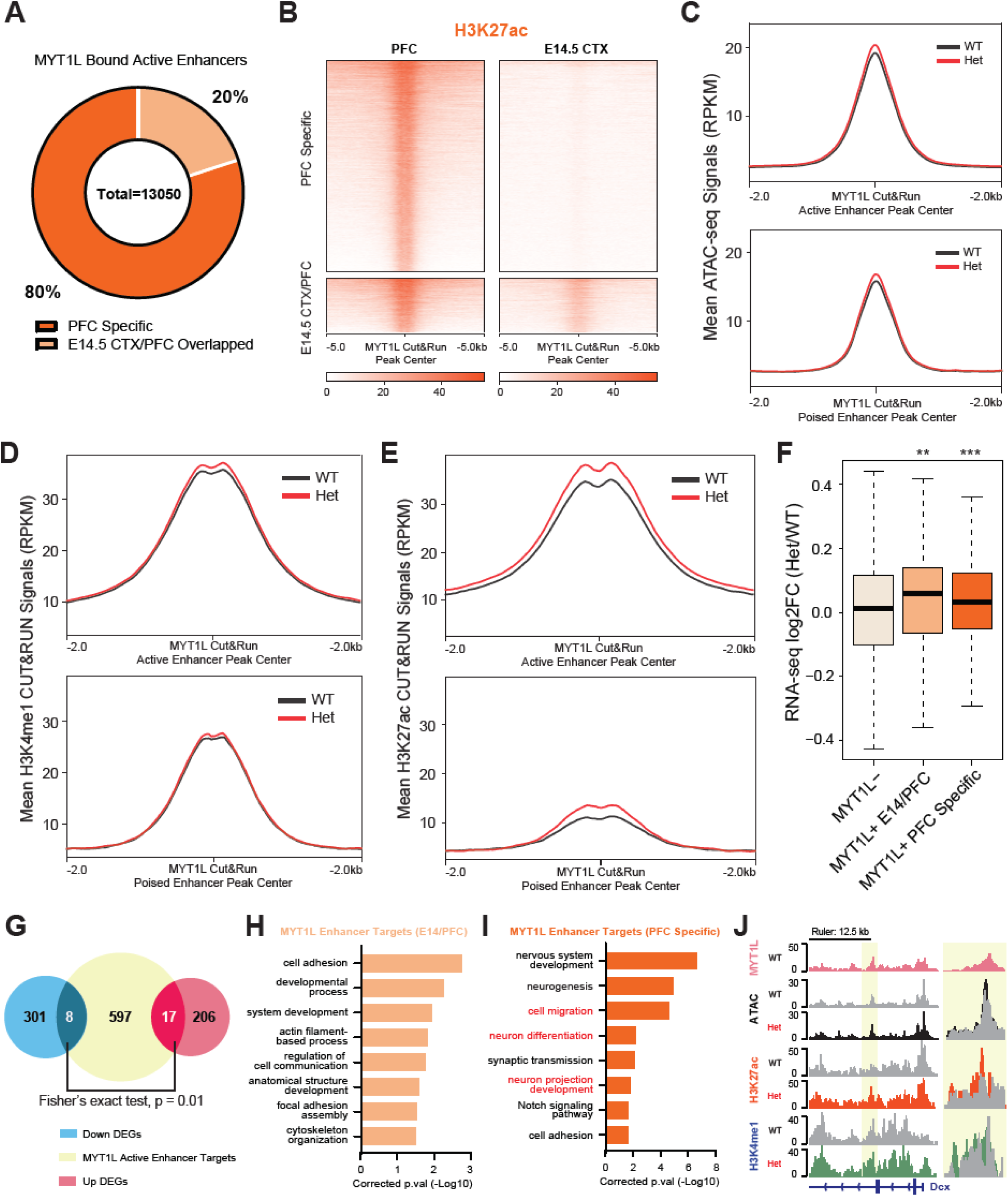
MYT1L suppresses enhancers that regulate neuronal migration and neuron projection development. (**A**) Majority of MYT1L bound active enhancers are PFC specific. (**B**) Heatmaps of MYT1L bound active enhancers in PFC. (**C**) MYT1L loss increases its bound active enhancers but not poised enhancers’ chromatin accessibility. (**D**) MYT1L loss increases H3K4me1 levels at MYT1L bound active enhancers but does not significantly increase H3K4me1 levels at MYT1L bound poised enhancers’. (**E**) MYT1L loss increases both its bound active and poised enhancers, H3K27ac levels. (**F**) MYT1L active enhancer target genes show increased expression in Het PFC. (**G**) Venn Diagram showing the overlaps among dDEGs, MYT1L active enhancer targets, and uDEGs, and there are more overlaps between MYT1L active enhancer targets and uDEGs than dDEGs (p = 0.01). (**H**) GO analysis on genes regulated by MYT1L bound E14 CTX/PFC overlapped active enhancers. (**I**) GO analysis on genes regulated by MYT1L bound PFC-specific active enhancers. (**J**) Representative genome browser track shows MYT1L-bound Dcx active enhancer has higher ATAC-seq signals, H3K4me1 and H3K27ac levels in Het PFC than WT. *p<0.05, **p<0.01, ***p<0.001, ****p<0.0001.

Next, we examined chromatin structure and histone landscape alterations at MYT1L+ enhancers in Het PFC to understand how MYT1L regulates enhancer activity. Unlike MYT1L+ promoters which showed no ATAC-differences, in Hets, the MYT1L+ active enhancers showed increased chromatin accessibility compared to WTs (**Fig 5C, S5B**), while MYT1L+ poised enhancers have no change (**Fig 5C, S5B**). When only looking at differential accessible regions (DARs, defined in (Chen et al. 2021)) annotated as active enhancers, up-regulated DARs have a higher percentage in MYT1L+ active enhancers than in MYT1L-active enhancers (**Fig S5C**). Likewise, DARs annotated as poised enhancers showed the same pattern (**Fig S5D**). This demonstrates that, at least on the chromatin accessibility, MYT1L tends to close the chromatin of its bound enhancers, and many DARs are direct effects. To better understand the enhancer activities, we looked into active histone marks, including H3K4me1 and H3K27ac. MYT1L active enhancer targets have increased H3K4me1 levels in Hets compared to WTs, while MYT1L poised enhancer targets showed unchanged H3K4me1 (**Fig 5D, S5E**). Notably, both MYT1L+ active and poised enhancers displayed increased enrichment of H3K27ac, a histone modification marking enhancer activation, in Het PFC (**Fig 5E,S5F**), suggesting MYT1L loss can also activate its bound poised enhancers. Together, these results indicate MYT1L normally facilitates repression of its bound enhancers, and MYT1L loss leads to aberrant enhancer activation, and more open chromatin.

Enhancers are important cis-regulatory elements for gene expression. Therefore, we again leveraged our RNA-seq datasets to understand how MYT1L together with enhancers control gene expression and to define the transcriptional consequences of MYT1L loss at enhancers. To find enhancer-gene-pairs (‘enhancer targets’, **Table S5**), we utilized EnhancerAtlas 2.0, a consensus enhancer prediction database based on multiple high throughput dataset including histone modifications, ATAC-seq, ChIA-seq, etc. (Gao and Qian 2020). We only focused on active enhancers since they are MYT1L’s major targets. Consistent with their increased active histone marks, MYT1L+ active enhancer targets tend to show increased gene expression in Het PFC compared with MYT1L-’s targets (**Fig 5F**). Likewise, when overlapping with DEGs, there are more uDEGs associated with MYT1L+ active enhancers compared to dDEGs, again emphasizing MYT1L’s primary role as a transcriptional repressor (**Fig 5G**).

We next examined the putative function of this enhancer regulation using GO, examining adult-specific MYT1L+ enhancers (**Fig. 5I**) and those enhancers bound at both ages (**Fig. 5H**). Since there are too few overlaps between MYT1L active enhancer targets and DEGs to perform GO (**Fig 5G**), we focused on all MYT1L+ active enhancer targets to obtain an overview of those enhancer functions. GO analysis on active enhancer targets found at both ages displayed enrichment of cytoskeleton pathways (**Fig 5H**). This is consistent with cytoskeleton related biological processes being constantly required from early development to adulthood. Meanwhile, PFC-specific active enhancer targets showed significant enrichment of neuronal migration and projection neuron development pathways (e.g., *Dcx*, **Fig 5I,J**), indicating the aberrant activation of earlier neuronal development programs in Het PFC seen at promoters can also be further exacerbated by dysregulation of MYT1L bound active enhancers.

## Discussion

Here, we utilized a MYT1L germline knockout mouse model to investigate MYT1L’s roles in neuronal maturation and the underlying molecular mechanisms. We first found that MYT1L loss increases the ratio of DL to UL neurons in the adult mouse cortex, consistent with the increase in the expression of known DL genes and decrease of UL genes observed in bulk RNAseq. To understand how MYT1L might mediate this shift in neuronal proportion, as well as to understand the consequences of MYT1L loss on both gene expression and epigenome marks, we mapped high confidence MYT1L binding targets as well as histone landscape using CUT&RUN on brain samples. Integrating CUT&RUN data with existing ATAC-seq and RNA-seq datasets, we identified that MYT1L directly bound a master regulator of DL fate, *Bcl11b* (Arlotta et al. 2005), and that this gene was upregulated when MYT1L was haploinsufficient in Het animals. We also saw a relative loss of the UL gene expression, though no loss of UL number. In addition, we detected widespread decrease in expression of later neuronal maturation genes, and sustained expression of early neuronal development programs. This finding defines a novel role for MYT1L in adult mouse brain to suppress early neuronal development programs by closing chromatin structures and erasing active histone markers at its binding sites, especially at enhancers. This sheds light on how MYT1L guides neuronal maturation and why MYT1L loss results in immature molecular and cellular signatures in the adult mouse brain.

MYT1L has been shown to repress non-neuronal genes to facilitate neuronal differentiation in MEFs reprogramming. Integrating MYT1L CUT&RUN and multi-omics data, we defined an additional role for MYT1L in repressing early neuronal development programs in the adult mouse cortex. We proposed that, MYT1L normally binds to both promoter and enhancer regions in postmitotic but immature neurons, recruits the SIN3B repressor complex containing HDAC2 and erases active histone marks to suppress earlier neuronal development genes (**Fig 6**) once this phase is complete. Shutting down earlier neuronal programs may allow postmitotic neurons to further mature to their final neuronal identities. Indeed, MYT1L Het PFC showed aberrant activation of promoters, enhancers, and thus gene expression associated with early neuronal development (**Fig 4, 5**), suggesting neurons are trapped in an immature stage. This epigenetic alteration may explain the disrupted transcriptional, morphological, and electrophysiological properties observed in this model (Chen et al. 2021).

**Figure 6:**
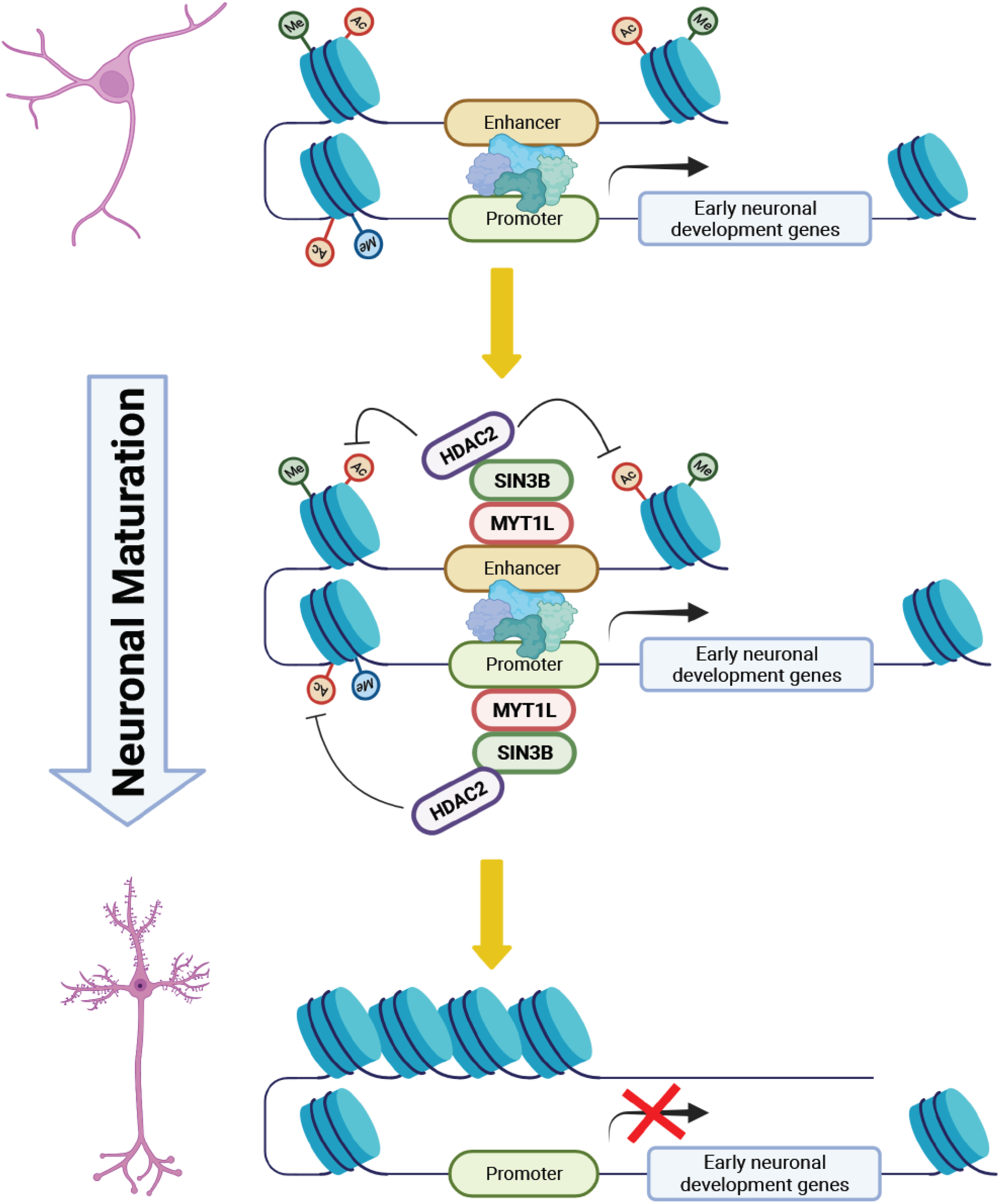
The model for MYT1L repressing early neuronal development programs to facilitate neuronal maturation.

In this study, CUT&RUN served as a powerful tool to profile MYT1L binding *in vivo*. While promoters remain frequent targets of MYT1L, we found 56% of MYT1L targets are enhancers, which is a much higher percentage than previously published ChIP-seq data (Mall et al. 2017). CUT&RUN is thought to have higher sensitivity and yield lower background signals compared with traditional ChIP-seq (Skene and Henikoff 2017). This may explain the larger number of MYT1L bound enhancers detected here than prior ChIP-seq. In addition, the MYT1L KO samples provided key control samples for optimizing the protocol, and were not available at the time of the ChIP-seq experiments. MYT1L peaks identified by CUT&RUN were not present in KOs, suggesting CUT&RUN with this antibody was very specific for MYT1L (**Fig 2C**). Consistent with this, the MYT1L core binding motif AAGTT has greater presence in peaks from CUT&RUN than ChIP-seq (76.4% vs 32.1%), though this can depend upon whether we use a very stringent analysis (using only the intersection of peaks found in 3 independent replicates) or a more sensitive approach of merging all samples into a single alignment file for peak calling, which may detect lower affinity binding sites. Meanwhile, CUT&RUN also exhibited different sensitivity between embryonic cortex and adult PFC, with many more MYT1L peaks were called from PFC (**Fig S2D,E**). Furthermore, peak enrichment was significantly improved in the PFC dataset than E14 cortex (**Fig S2A**). Although most of E14 peaks from both peak calling methods can be recovered in PFC experiments (**Fig S2C,E**), it is hard to tell if the unique peaks from two developmental time points are derived from differential bindings of MYT1L at the different ages, or rather a CUT&RUN sensitivity difference. CUT&RUN on more homogenous cell populations or single-cell TF-DNA interaction profiling assays (Cammack et al. 2020; Moudgil et al. 2020) will be needed to differentiate these two possibilities in the future.

MYT1L is a pro-neuronal TF that has been shown to regulate neuronal differentiation in many *in vitro* studies (Heavner et al. 2020; Mall et al. 2017; Vierbuchen et al. 2010). Here, utilizing MYT1L germline knockout mice, we explored long-term consequences of MYT1L loss on neuronal molecular maturation *in vivo*. We found MYT1L levels influence neurons’ layer specific identity in the adult brain, while MYT1L mutation leads to increased DL neuron numbers and an up-regulation of the corresponding genes. This is consistent with the phenotype from a MYT1L shRNA knockdown study on primary cortical neuron cultures (Heavner et al. 2020). Another MYT1L shRNA knockdown experiment by *in utero* electroporation also showed that neurons with MYT1L loss fail to migrate into upper cortical layers during mouse embryonic development (Mall et al. 2017), which might eventually result in increased DL neuron numbers in the adult brain. CUT&RUN experiments revealed that MYT1L directly binds to genomic regions associated with early neuronal development, including neuronal migration and projection development, as well as the DL gene *Bcl11b*. Combining RNA-seq and ATAC-seq datasets, we propose a role for MYT1L in providing proper suppression of these earlier neuronal development programs (ENDPs). We believe this can explain how MYT1L loss can simultaneously lead to precocious neuronal differentiation in embryos (and thus later microcephaly), yet prolonged neuronal immaturity in adults. Specifically, in embryos, MYT1L could normally help prevent a too-early expression of ENDPs. Thus in mutant embryos, there is a bias in the progenitors to move too quickly from proliferation into early neuronal differentiation. Yet, once the animals pass the age at which these EDNPs are needed at a high level, they are also unable to recruit sufficient SIN3B and downstream HDACs to de-acetylate many of these promoters and enhancers, and thus turn down the EDNPs to allow the neurons to complete their maturation. It is also possible that in addition to any effects of BCL11B on DL number directly, increased DL neurons gene expression pattern and decreased UL pattern might represent another aspect of the precocious differentiation and immaturity since DL neurons belong to an earlier neurodevelopmental trajectory than UL neurons in the cortex, which forms in an in-side-out pattern during development (Shepherd and Rowe 2017).

Despite consensus findings on MYT1L’s role in facilitating neuronal differentiation both *in vitro* and now *in vivo*, how MYT1L does so remains poorly understood. Unlike the repressive effects of MYT1L overexpression on non-neuronal genes described in transdifferentiation system (Mall et al. 2017), we saw no obvious activation of non-neuronal genes upon MYT1L loss (Chen et al. 2021). Likewise, we observed no obvious MYT1L binding at promoters of non-neural genes (e.g., liver, fibroblasts) in the CUT&RUN *in vivo*, in contrast to the ChIP-seq data in trans differentiating cells. Identical effects on different targets between *in vitro* and *in vivo* systems indicate MYT1L’s repressive functions are context dependent, and ectopic expression of MYT1L might change its functions from those under physiological conditions.

In addition to functioning as a transcriptional repressor, evidence has been accumulated to suggest MYT1L can also activate transcription. We found MYT1L loss decreases its bound promoters’ accessibility via ATAC-seq when defining targeting using ChIP-seq *in vitro* (Mall et al. 2017; Chen et al. 2021). And while on average the DARs and DEGs at MYT1L bound genes tended to indicate it more often a repressor in WT brain, many individual loci showed a response in mutants more consistent with a loss of an activator (**Fig 4F-G, 5F-G, S4C-D**). Meanwhile, the N-terminus of MYT1L alone is sufficient to activate transcription in the luciferase reporter assay *in vitro* (Manukyan et al. 2018). To explain these two faces of MYT1L, a “ready-set-go” model has been proposed, where MYT1L co-operates with different co-factors to control neuronal gene transcription (Chen et al. 2022). Our study identified several co-factor candidates for MYT1L *in vivo*, including both transcriptional activators (SP1 and ELK1) and repressors (SIN3B), providing important hints for developing future models of how MYT1L tunes neuronal gene expression at different gene classes. High throughput techniques, including massive parallel reporter assays (MPRA) (Mulvey et al. 2021), can be leveraged in the future to further examine the motif and cofactor requirements at MYT1L targets for repression and activation respectively.

We also noticed that not all targets bound by MYT1L responded with a uniform magnitude to MYT1L heterozygosity, and this may shed some light on the phenotype in MYT1L Het mice and haploinsufficient patients. Determining why certain MYT1L-bound genes are specifically sensitive to MYT1L levels will be another important future direction. It is interesting to note that neurite outgrowth genes, which are often disrupted, may be more dependent on neural activity dependent gene expression than other processes. Given MYT1L is often cobinding with activity dependent genes like Fos and Jun at enhancers, it may be that MYT1L is needed to turn off activity-dependent signals like these after their activation. If so, this would fit with an earlier theory suggesting IDD/ASD may be a general consequence of mistimed activity dependent gene expression (Ebert and Greenberg 2013), though this would require further manipulations to assess in MYT1L syndrome models.

Collectively, we mapped MYT1L binding targets via CUT&RUN and defined a function in suppressing EDNPs in the adult mouse brain. In addition, the data provided here should provide a foundation to study how co-binding partners and or neural activity might influence the function of MYT1L in gene expression and histone modification. Such detailed investigations on MYT1L functions both in WT and MYT1L Syndrome mouse models could advance our understanding of the complicated progression of neuronal development programs in both physiological and pathological conditions.

## Methods

### Animal models

All procedures using mice were approved by the Institutional Care and Use Committee at Washington University School of Medicine. All mice used in this study were bred and maintained in the vivarium at Washington University in St. Louis in individually ventilated (36.2 x 17.1 x 13 cm) or static (28.5 x 17.5 x 12 cm; post-weaning behavior only) translucent plastic cages with corncob bedding and *ad libitum* access to standard lab diet and water. Animals were kept at 12/12 hour light/dark cycle, and room temperature (20-22°C) and relative humidity (50%) were controlled automatically. For all experiments, adequate measures were taken to minimize any pain or discomfort. Breeding pairs for experimental cohorts comprised *Myt1l* Hets and wild type C57BL/6J mice (JAX Stock No. 000664) to generate male and female *Myt1l* Het and WT littermates. For embryonic CUT&RUN, *Myt1l* Het x Het breeding pairs were used to generate *Myt1l* WT and homozygous mutant littermates. Animals were weaned at P21, and group-housed by sex and genotype, until tissue harvest at P60 of age. Biological replicates for all experiments were sex and genotype balanced.

### Gene Set Enrichment Analysis (GSEA)

GSEA was performed as described before (Subramanian et al. 2005) using GSEA v4.2.3 (https://www.gsea-msigdb.org/gsea/index.jsp). Deep layer neuron and upper layer neuron gene lists were obtained from Heavner et al., 2020. RNA-seq datasets on MYT1L germline knockout mice were obtained from (Chen et al. 2021). All analysis was performed with “gene_set” as permutation type and 1,000 permutations. Significant enrichment was determined by FDR < 0.1 cut-off.

### Histopathology

Mice (5 WTs and 5 Hets for BRN2 staining, 7 WTs and 6 Hets for CTIP2 staining, sex balanced, at the age of P60) were deeply anesthetized and transcardially perfused with 4% paraformaldehyde in PBS. Whole brains were weighed and serially sectioned in the coronal plane at 75 μm using a vibratome and immunolabeled for either CTIP2 (a marker for cortical layers V/VI) or BRN2 (a marker for cortical layers II-IV). For each antibody, a set consisting of every eighth section was isolated and slide mounted. After drying overnight, antigen retrieval was performed by immersing in citrate buffer (pH 6.0) and pressure cooking for 10 minutes. The slides were then quenched in 3% hydrogen peroxide in absolute methanol for 10 minutes, immersed for 1 hour in a blocking solution (2% bovine serum albumin, 0.2% dry milk, 0.8% TX-100 in PBS), and incubated overnight with a 1:500 dilution of either Ctip2 (CAT#ab18465; Abcam, Burlington, MA) or Brn2 (CAT#sc-393324; Santa Cruz, Dallas, TX). The next morning, Ctip2 or Brn2 incubated sections were reacted with appropriate biotinylated secondary antibody for 1 hour (B7139; Sigma-Aldrich, St. Louis, MO; 1:200 or BA-9200; Vector Labs, Burlingame, CA; 1:200 respectively). The sections were then reacted with an avidin-biotin conjugate (ABC kit) for 1 hour and visualized using the chromogen VIP (Vectastatin Elite ABC kit and Vector VIP kits; Vector Labs, Burlingame, CA).

### Stereology

After immunolabeling, Ctip2 or Brn2 positive neurons were stereologically quantified using Stereoinvestigator Software (v 2019.1.3, MBF Bioscience, Williston, Vermont, USA) running on a Dell Precision Tower 5810 computer connected to a QImaging 2000R camera and a Labophot-2 Nikon microscope with electronically driven motorized stage. A rater, blind to treatment, stereologically quantified the number of positively stained cells using the unbiased optical fractionator method. To restrict counting to cortical regions with six layers, cell counts were performed on sections where the corpus callosum was visible and only in the neocortex (this excludes the allocortex, piriform, entorhinal, and retrosplenial cortices). Since each antibody labels specific cortical layers, volumes were calculated for only layers V-VI for Ctip2 and I-IV for Brn2 (Brn2 did label cells in layers V-VI but these were not counted). Finally, a density was calculated by dividing the total number of positive cells by total volume for each antibody. Since maternal care, litter size, and other factors can cause litter effects, data were normalized by dividing each value by the average of the wild-type animals within each litter.

### CUT&RUN on embryonic and adult prefrontal cortex

CUT&RUN was performed on the embryonic and adult prefrontal cortex as previously described (Skene and Henikoff 2017; Brodie-Kommit et al. 2021). Three biological replicates were included for each age and genotype. Briefly, E14 mouse embryonic cortex or P60 mouse prefrontal cortex were dissected out, and nuclei were isolated using Nuclei EZ Prep Buffer (Sigma 4432370) and counted on cell cytometer. 300k nuclei were bound to the Concanavalin A-coated beads for each CUT&RUN reaction. Then, each aliquot of bead/nuclei were incubated with a primary antibody, including Rb-MYT1L (0.5 μg), Rb-H3K4me1 (1 μg), Rb-H3K4me3 (1 μg), Rb-H3K27ac (1 μg), and Rb IgG (1 μg), at 4 °C on the nutator for overnight. Next, to bind pAG-MNase fusion protein to the antibodies, beads were incubated with diluted CUTANA pAG-MNase (1:20) on the rotator at 4 °C for 1 hour. Chromatin digestion was performed at 0 °C with the addition of CaCl_2_ (100 mM) for 30min. To digest the RNA and release the cleaved DNA fragments, reactions were incubated with Stop Buffer at 37 °C for 30min in the thermocycler. Magnetic stands were used to bind beads afterwards, and supernants containing DNA fragments were retrieved for sequencing library preparation.

### CUT&RUN library preparation and the next generation sequencing

DNA fragments were extracted from CUT&RUN experimental supernatants by Phenol/Chloroform/isoamyl Alcohol (pH 7.9) mix (Skene and Henikoff 2017). KAPA HyperPrep Kit (KK8504) was used to generate dual-indexed sequencing libraries. Generated libraries were then purified using Mag-Bind beads. Finally, a robust nucleosome peaks pattern was confirmed as a quality control using an Agilent Tapestation and HS D1000 tapes. Finally, libraries were submitted to GTAC@MGI at Washington University School of Medicine for Illumina sequencing using a Novaseq instrument, with a targeted read depth of 50M reads per MYT1L and IgG library and 10M reads per histone library.

### CUT&RUN data analysis

Raw reads were trimmed by Trimmomatic software to remove adapter sequence. Fastqc was used to check read quality before and after trimming. Then reads were mapped to the mm10 genome by Bowtie2. Mitochondrial reads (Samtools), PCR duplicates (Picard), non-unique alignments (MAPQ > 30), and unmapped reads (Samtools) were filtered out. MYT1L peaks were called from both individual biological replicates (q<0.05) as well as from merged bam files (merged by genotype, q<0.01) MACS2 using IgG as background. Histones’ peaks were called from merged bam files by MACS2 (q<0.05) using down-sampled IgG as background. With MYT1L peaks called from biological replicates, bedTools was used to find intersecting peaks among 3 replicates. With MYT1L peaks called from merged bam files, bedTools was also utilized to exclude the 208peaks found in KO samples. Next, peaks were annotated by Homer and then grouped into subcategories as described in the next section. Peak heatmaps were generated by the bedTools plotHeatmap function. Genome track graphs were generated using Integrated Genomics Viewer (https://igv.org/). In order to compare changes of histone levels between WT and Het PFC, read counts within MYT1L CUT&RUN peaks were derived from individual histone CUT&RUN bam file using bedtools. Then, read counts were normalized to corresponding library sequencing depth and the average coverage was calculated from biological replicates within the genotype.

### Definition of active and poised enhancers

Active and poised enhancers were defined as previously described (Creyghton et al. 2010). Briefly, enhancers were defined as H3K4me1 peaks located outside of the promoter regions (TSS ±1kb), with an absence of H3K4me3. Enhancers that overlap with H3K27ac peaks were categorized as active enhancers, and those without H3K27ac were defined as poised enhancers.

### Motif analysis

De novo motif discovery was performed using both Homer and monaLisa. For Homer usage, full length peaks were fed into the software, and ATAC-seq peaks from the same brain region and the same age were used as background. For monaLisa usage, promoter, active enhancer, and poised enhancer targets were grouped into separate bins and tested individually. To avoid length bias, peaks were resized into fixed-size regions around the peak midpoint before running the analysis. Then, the k-mer enrichment analysis was performed using monaLisa to examine the MYT1L core binding motif enrichment using 5 as unbiased motif length and ATAC-seq peaks as background. Finally, known motif finding was performed using monaLisa and the JASPAR2020 motif database on MYT1L bound peaks. plotMotifHeatmaps was used to visualize significantly enriched known motifs with FDR<1e-5, and with TFs showing expression in PFC RNA-seq dataset were plotted in the heatmap (**Figure 4A**). To validate the motif analysis, different TFs’ ChIP-seq data from most relevant tissue types were acquired from the ChIP-Atlas (https://chip-atlas.org/) (Oki et al. 2018). Specifically, SP1 ChIP-seq on striatal neurons, ELK1 ChIP-seq on striatal neurons, JUNB ChIP-seq on activated cortical neurons, MEF2A ChIP-seq on PFC, NEUROD1 ChIP-seq on striatal neurons, and NEUROD2 ChIP-seq on striatal neurons cortex were fed into ChIPpeakAnno Package to overlap with MYT1L CUT&RUN data with a maximum gap of 200bp.

### Predicting enhancer-gene-pairs

Enhnacer-gene-pairs were predicted using EnhancerAtlas 2.0 (http://www.enhanceratlas.org/). Specifically, E14.5 Brain, Brain, and Neuron database were selected to annotate different subgroups of enhancers with their putative targeting genes.

### Gene Ontology (GO) analysis

GO analysis was performed using the BiNGO app in Cytoscape. *p* values were adjusted by Benjamini-Hochberg FDR correction, and FDR < 0.05 cut-off was used to determine significant enrichments. Full GO analysis results can be seen in **Table S4**.

### Integration between CUT&RUN dataset and ATAC-seq

ATAC-seq datasets, including peak files and differential accessible region analysis results, were obtained from (Chen et al. 2021). The ChIPpeakAnno package was used to find overlapping peaks among MYT1L CUT&RUN, histones’ CUT&RUN, and ATAC-seq datasets. The maxgap value was set as 0. Then, fold change values for histone enrichments and ATAC-seq signals were retrieved for MYT1L bound and unbound regions respectively. Boxplots were generated using R built-in functions.

### Co-Immunoprecipitation

WT C57BL/6J P1 mouse cortex was dissected out in cold PBS and put into the lysis buffer (150mM NaCl, 50mM Tris, 1% Triton-X) with protease inhibitors for homogenization. Brain lysates were centrifuged at 15,000 X g for 10 min at 4°C. Then, supernatants were pre-cleared with Protein A or G Dynabeads® for 1 hour at 4°C. Pre-cleared lysates were used as immunoprecipitate inputs. To bind antibodies to beads, 1-5 μg rabbit anti-MYT1L antibody was added into a 200 μL lysis buffer with 20 μL Dynabeads® and rotated for 20 min at RT. The bead-antibody complex was washed twice with lysis buffer and then incubated with 100 μL of pre-cleared brain lysates rotated at 4°C overnight. The bead-antibody-antigen complex then was washed four times with lysis buffer and resuspended in 15 μL lysis buffer plus 15 μL 2X sample buffer.The mixture was boiled for 10 min, and supernatants were separated from beads using a magnetic stand. Supernatants were then subjected to immunoblotting experiments for detecting candidate co-factors as described (Chen et al. 2021).

### Statistical analysis

Statistical analyses and data graphing were performed using GraphPad Prism (v.8.2.1), and R(v.4.0.0). Prior to analyses, data was screened for missing values and for the fit of distributions with assumptions underlying univariate analysis. Means and standard errors were computed for each measure. Analysis of variance (ANOVA), including repeated measures or mixed models, was used to analyze data where appropriate. One-sample *t*-tests were used to determine differences from chance. For data that did not fit univariate assumptions, non-parametric tests were used or transformations were applied. Fisher’s exact tests were used to assess MYT1L bound and unbound DAR distributions. Mann–Whitney U test was used to examine gene expression, ATAC-seq signal, and histone enrichment differences among groups. Multiple pairwise comparisons were subjected to Bonferroni correction or Dunnett correction. Figure schematics were generated using BioRender. All statistical data can be found in **Table S6**.

## Data availability

The codes for analyzing Illumina sequencing, ATAC-seq, and RNA-seq generated in this study are available via Bitbucket: ???. The CUT&RUN raw reads as well as counts data are available at GEO with reference ID GSE???. Any additional information required to reanalyze the data reported in this work paper is available from the Lead Contact upon request.

## Supplemental Information

**Table S1. CUT&RUN sequencing metrics and peak profiles**

**Table S2. Subcategories of MYT1L Targets**

**Table S3. Enhancer annotations**

**Table S4. Gene Ontology analysis on MYT1L Targets**

**Table S5. MYT1L promoter and enhancer target gene predictions**

**Table S6. Statistical analysis results**

## Author Contributions

Conceptualization: J.C., J.D.D.; Methodology: J.C., K.N., J.D.D.; Formal Analysis: J.C., N.F., K.N.; Investigation: J.C., N.F., K.N.; Data Curation: J.C., N.F., K.N.; Writing – Original Draft: J.C., N.F., K.N., J.D.D.; Writing – Review & Editing: J.C., N.F., K.N., J.D.D.; Visualization: J.C., J.C., N.F., K.N.; Supervision: K.N., J.D.D.; Project Administration: J.C., J.D.D.; Funding Acquisition: J.C., J.D.D..

## Acknowledgements

We thank Dr. Brian Clark, Dr. Harrison Gabel, Dr. Kristen Kroll, Florian Colin, Nicole Hamagami, Diana Christian for technical assistance and scientific advice. Funding was provided by The Jakob Gene Fund, the Mallinckrodt Institute of Radiology at Washington University School of Medicine, McDonnell International Scholars Academy (J.C.), and the NIH: (R01MH107515, R01MH124808 to JDD, and NIH 5UL1TR002345 (ICTS) and P50 HD103525 (IDDRC)).

---

**Supplemental Figure 1:**
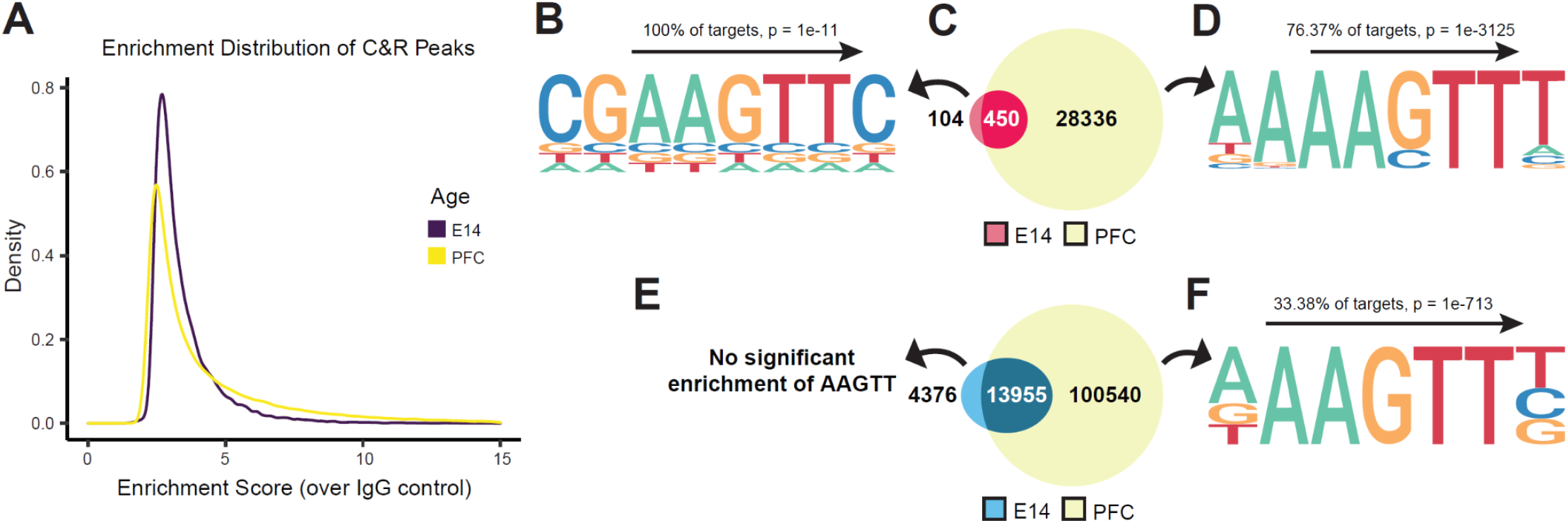
CUT&RUN is more efficient for MYT1L binding profiling on P60 PFC than on E14 CTX. (**A**) MYT1L CUT&RUN peaks called from the merged PFC alignment file showed decreased density of low enrichment peaks but increased density of high enrichment peaks compared to E14 CUT&RUN peaks. (**B**) Homer de novo motif finding shows significant enrichment of MYT1L core binding motif AAGTT in E14 CUT&RUN intersected peaks. (**C**) Overlaps between MYT1L PFC and E14 intersected CUT&RUN peaks. (**D**) Homer de novo motif finding shows significant enrichment of MYT1L core binding motif AAGTT in PFC CUT&RUN intersected peaks. (**E**) Overlaps between MYT1L PFC and E14 CTX CUT&RUN peaks called from the merged alignment file. (**F**) Homer de novo motif finding shows significant enrichment of MYT1L core binding motif AAGTT in PFC CUT&RUN peaks called from the merged alignment file.

**Supplemental Figure 2:**
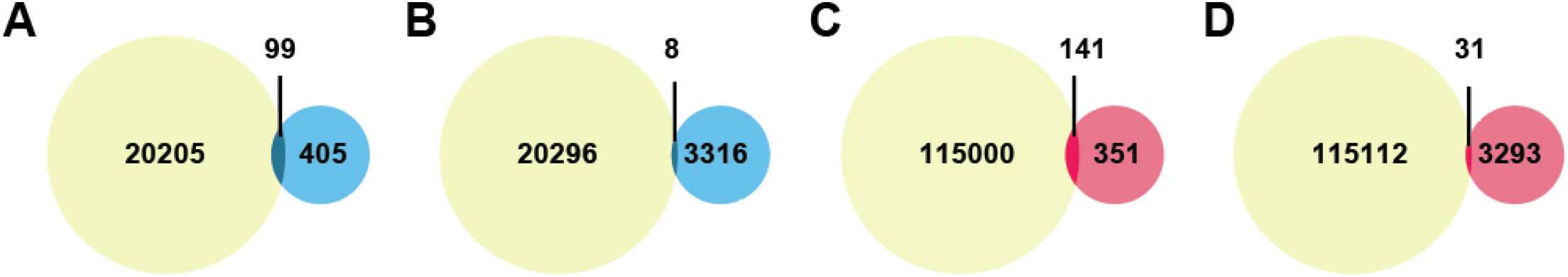
(**A**) No significant overlap was found between MYT1L E14 CTX CUT&RUN peaks called from the merged alignment file and MYT1L ChIP-seq peaks from the mouse E14 brain. (**B**) No significant overlap was found between MYT1L E14 CTX CUT&RUN peaks called from the merged alignment file and MYT1L ChIP-seq peaks from MEFs overexpressing MYT1L, BRN2, and ASCL1. (**C**) No significant overlap was found between MYT1L PFC CUT&RUN peaks called from the merged alignment file and MYT1L ChIP-seq peaks from the mouse E14 brain. (**D**) No significant overlap was found between MYT1L PFC CUT&RUN peaks called from the merged alignment file and MYT1L ChIP-seq peaks from MEFs overexpressing MYT1L, BRN2, and ASCL1. Color code: CUT7RUN (yellow), MEFs ChIP-seq (blue), and E14 brain ChIP-seq (red).

**Supplemental Figure 3:**
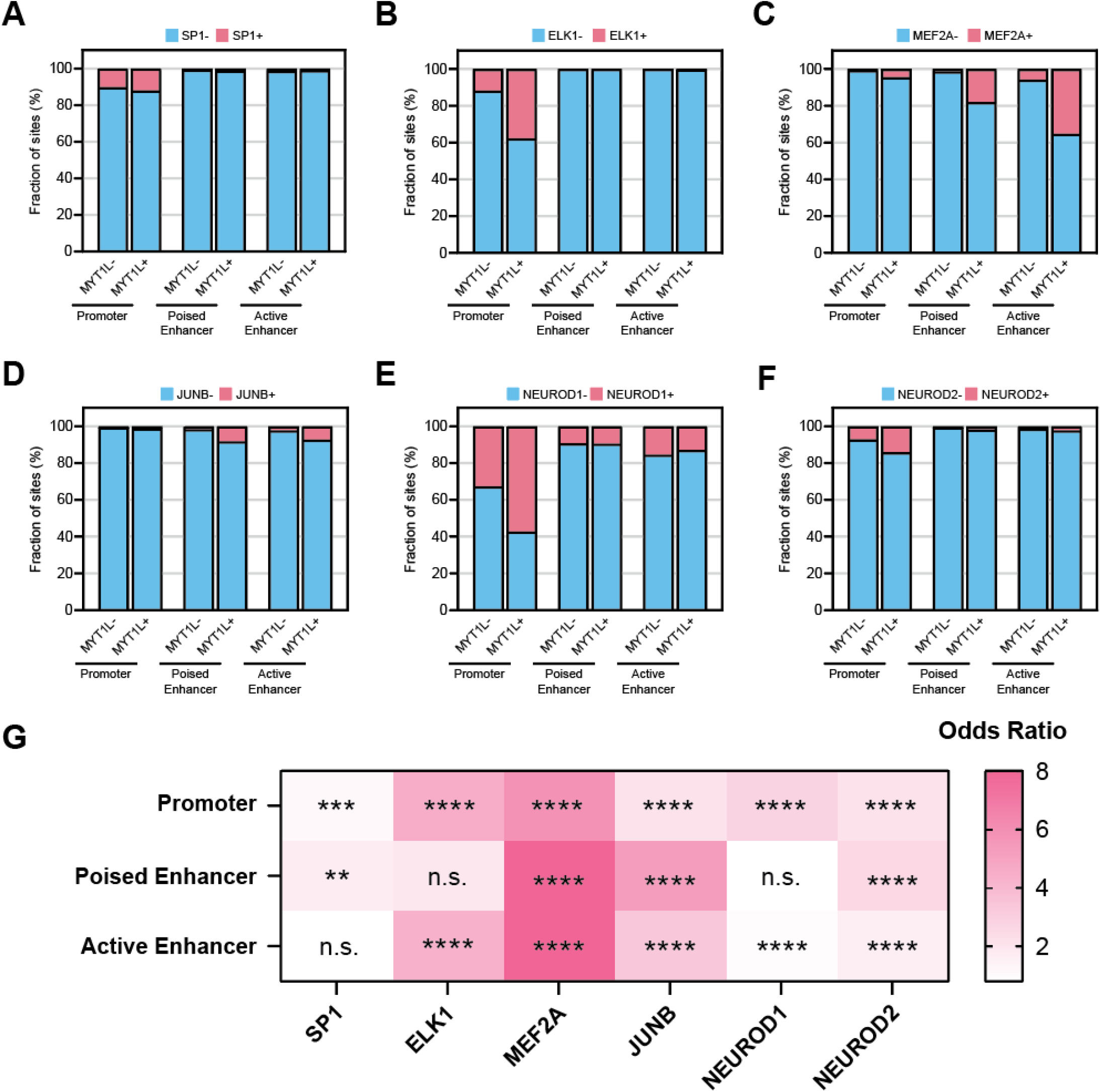
MYT1L co-occupies with a different set of transcriptional factors at promoters and enhancers. (**A**) MYT1L+ promoters have a significantly higher percentage of SP1 binding compared to MYT1L-promoters, while no obvious difference was observed in enhancers. The same pattern was observed for (**B**) ELK1 as well. (**C**) MYT1L+ enhancers have a significantly higher percentage of MEF2A binding compared to MYT1L-promoters, while no big difference was observed in promoters. The same pattern was observed for (**D**) JUNB as well. (**E**) MYT1L+ promoters have a significantly higher percentage of NEUROD1 binding compared to MYT1L-promoters, while no obvious difference was observed in enhancers. The same pattern was observed for **(F)** NEUROD2 as well. (**G**) Fisher’s exact test showed different TFs’ enrichment in MYT1L+ over MYT1L-genomic regions. *p<0.05, **p<0.01, ***p<0.001, ****p<0.0001.

**Supplemental Figure 4:**
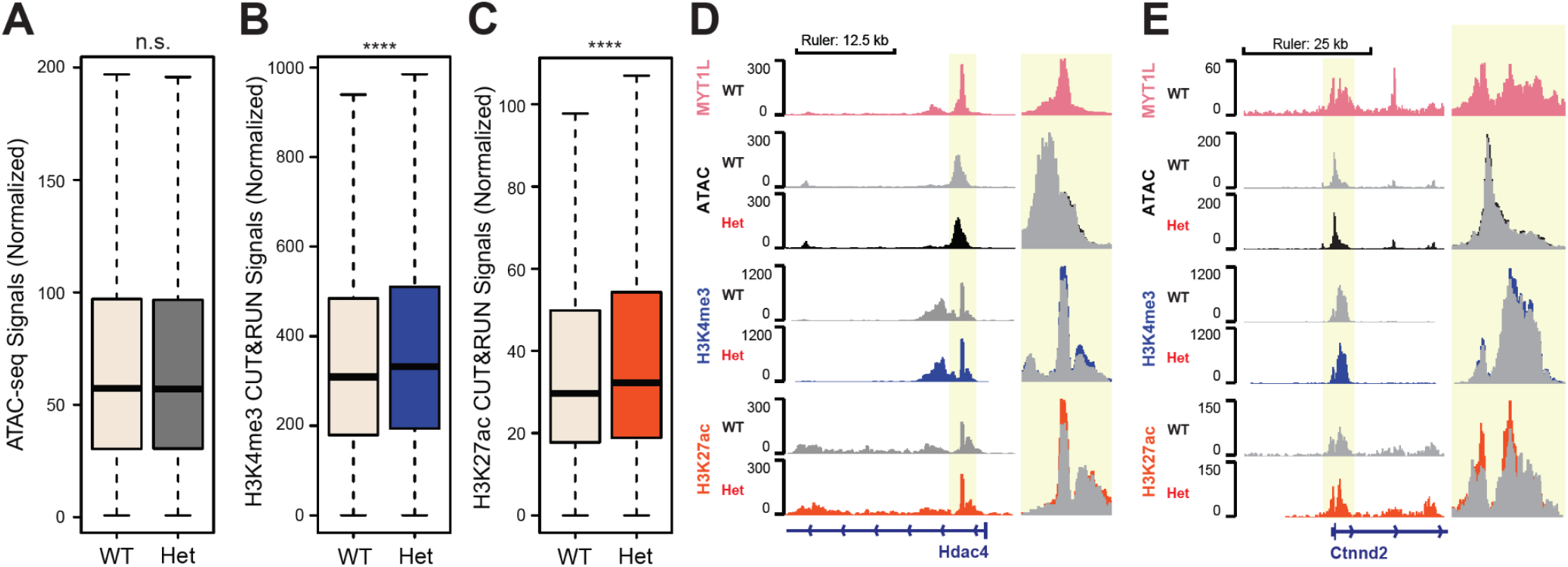
MYT1L loss increases H3K4me3 and H3K27ac levels at promoters. **(A)** MYT1L loss did not alter its bound promoters’ chromatin accessibility. **(B)** MYT1L Het PFC has higher levels of H3K4me3 at MYT1L promoter targets compared to WTs. **(C)** MYT1L Het PFC has higher levels of H3K27ac at MYT1L promoter targets compared to WTs. (**D**) Representative genome browser track of MYT1L bound Hdac4 promoter. (**E**) Representative genome browser track of MYT1L bound Ctnnd2 promoter. *p<0.05, **p<0.01, ***p<0.001, ****p<0.0001.

**Supplemental Figure 5:**
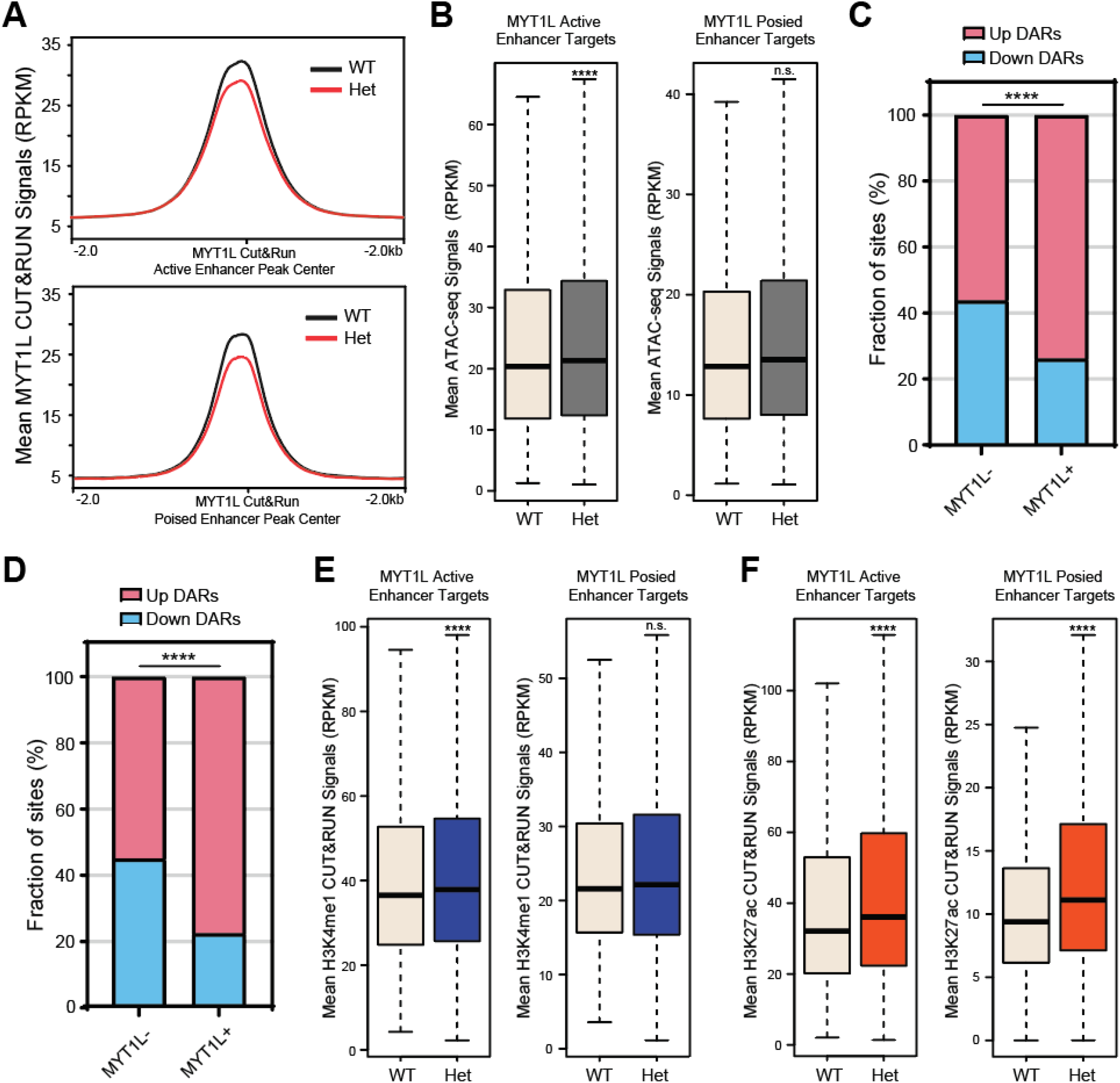
MYT1L loss increases activation marks of its bound enhancers. **(A)** MYT1L Het PFC has reduced MYT1L binding at active (top) and poised (bottom) enhancers. (**B**) MYT1L loss increases its bound active (left) but not poised (right) enhancers’ chromatin accessibility. (**C**) MYT1L bound active enhancer DARs have a higher percentage of DARs that increase accessibility than MYT1L unbound active enhancer DARs. (**C**) MYT1L bound poised enhancer DARs have a higher percentage of DARs that increase accessibility than MYT1L unbound poised enhancer DARs. **(E)** MYT1L Het PFC has higher levels of H3K4me1 at both active (left) and poised (right) enhancer targets compared to WTs. **(F)** MYT1L Het PFC has higher levels of H3K27ac at both active (left) and poised (right) enhancer targets compared to WTs. *p<0.05, **p<0.01, ***p<0.001, ****p<0.0001.

## Reference

Almazan G, Lefebvre DL, Zingg HH. 1989. Ontogeny of hypothalamic vasopressin, oxytocin and somatostatin gene expression. Developmental Brain Research 45: 69–75.

Arlotta P, Molyneaux BJ, Chen J, Inoue J, Kominami R, Macklis JD. 2005. Neuronal Subtype-Specific Genes that Control Corticospinal Motor Neuron Development In Vivo. Neuron 45: 207–221.

Arlotta P, Molyneaux BJ, Jabaudon D, Yoshida Y, Macklis JD. 2008. Ctip2 Controls the Differentiation of Medium Spiny Neurons and the Establishment of the Cellular Architecture of the Striatum. J Neurosci 28: 622–632.

Bainor AJ, Saini S, Calderon A, Casado-Polanco R, Giner-Ramirez B, Moncada C, Cantor DJ, Ernlund A, Litovchick L, David G. 2018. The HDAC-Associated Sin3B Protein Represses DREAM Complex Targets and Cooperates with APC/C to Promote Quiescence. Cell Reports 25: 2797–2807.e8.

Bedogni F, Hodge RD, Elsen GE, Nelson BR, Daza RAM, Beyer RP, Bammler TK, Rubenstein JLR, Hevner RF. 2010. Tbr1 regulates regional and laminar identity of postmitotic neurons in developing neocortex. Proceedings of the National Academy of Sciences 107: 13129–13134.

Black JC, Van Rechem C, Whetstine JR. 2012. Histone Lysine Methylation Dynamics: Establishment, Regulation, and Biological Impact. Molecular Cell 48: 491–507.

Blanchet P, Bebin M, Bruet S, Cooper GM, Thompson ML, Duban-Bedu B, Gerard B, Piton A, Suckno S, Deshpande C, et al. 2017. MYT1L mutations cause intellectual disability and variable obesity by dysregulating gene expression and development of the neuroendocrine hypothalamus ed. Z. Stark. PLoS Genet 13: e1006957.

Brodie-Kommit J, Clark BS, Shi Q, Shiau F, Kim DW, Langel J, Sheely C, Ruzycki PA, Fries M, Javed A, et al. 2021. Atoh7-independent specification of retinal ganglion cell identity. Science Advances 7: eabe4983.

Cammack AJ, Moudgil A, Chen J, Vasek MJ, Shabsovich M, McCullough K, Yen A, Lagunas T, Maloney SE, He J, et al. 2020. A viral toolkit for recording transcription factor–DNA interactions in live mouse tissues. PNAS 117: 10003–10014.

Campbell K. 2005. Cortical Neuron Specification: It Has Its Time and Place. Neuron 46: 373–376.

Carullo NVN, Day JJ. 2019. Genomic Enhancers in Brain Health and Disease. Genes 10: 43.

Chen J, Lambo ME, Ge X, Dearborn JT, Liu Y, McCullough KB, Swift RG, Tabachnick DR, Tian L, Noguchi K, et al. 2021. A MYT1L syndrome mouse model recapitulates patient phenotypes and reveals altered brain development due to disrupted neuronal maturation. Neuron. https://www.sciencedirect.com/science/article/pii/S0896627321006814 (Accessed October 18, 2021).

Chen J, Yen A, Florian CP, Dougherty JD. 2022. MYT1L in the making: emerging insights on functions of a neurodevelopmental disorder gene. Transl Psychiatry 12: 1–8.

Coursimault J, Guerrot A-M, Morrow MM, Schramm C, Zamora FM, Shanmugham A, Liu S, Zou F, Bilan F, Le Guyader G, et al. 2021. MYT1L-associated neurodevelopmental disorder: description of 40 new cases and literature review of clinical and molecular aspects. Hum Genet. https://doi.org/10.1007/s00439-021-02383-z (Accessed December 1, 2021).

Creyghton MP, Cheng AW, Welstead GG, Kooistra T, Carey BW, Steine EJ, Hanna J, Lodato MA, Frampton GM, Sharp PA, et al. 2010. Histone H3K27ac separates active from poised enhancers and predicts developmental state. Proceedings of the National Academy of Sciences 107: 21931–21936.

Dixit R, Wilkinson G, Cancino GI, Shaker T, Adnani L, Li S, Dennis D, Kurrasch D, Chan JA, Olson EC, et al. 2014. Neurog1 and Neurog2 Control Two Waves of Neuronal Differentiation in the Piriform Cortex. J Neurosci 34: 539–553.

Ebert DH, Greenberg ME. 2013. Activity-dependent neuronal signalling and autism spectrum disorder. Nature 493: 327–337.

Gao T, Qian J. 2020. EnhancerAtlas 2.0: an updated resource with enhancer annotation in 586 tissue/cell types across nine species. Nucleic Acids Research 48: D58–D64.

Götz M, Huttner WB. 2005. The cell biology of neurogenesis. Nat Rev Mol Cell Biol 6: 777–788.

Hayakawa T, Ohtani Y, Hayakawa N, Shinmyozu K, Saito M, Ishikawa F, Nakayama J. 2007. RBP2 is an MRG15 complex component and down-regulates intragenic histone H3 lysine 4 methylation. Genes to Cells 12: 811–826.

Heavner WE, Ji S, Notwell JH, Dyer ES, Tseng AM, Birgmeier J, Yoo B, Bejerano G, McConnell SK. 2020. Transcription factor expression defines subclasses of developing projection neurons highly similar to single-cell RNA-seq subtypes. Proceedings of the National Academy of Sciences 117: 25074–25084.

Heintzman ND, Hon GC, Hawkins RD, Kheradpour P, Stark A, Harp LF, Ye Z, Lee LK, Stuart RK, Ching CW, et al. 2009. Histone modifications at human enhancers reflect global cell-type-specific gene expression. Nature 459: 108–112.

Jiang Y, Yu VC, Buchholz F, O’Connell S, Rhodes SJ, Candeloro C, Xia Y-R, Lusis AJ, Rosenfeld MG. 1996. A Novel Family of Cys-Cys, His-Cys Zinc Finger Transcription Factors Expressed in Developing Nervous System and Pituitary Gland. Journal of Biological Chemistry 271: 10723–10730.

Kim S, Oh H, Choi SH, Yoo Y-E, Noh YW, Cho Y, Im GH, Lee C, Oh Y, Yang E, et al. 2022. Postnatal age-differential ASD-like transcriptomic, synaptic, and behavioral deficits in Myt1l-mutant mice. Cell Reports 40: 111398.

Kroon T, van Hugte E, van Linge L, Mansvelder HD, Meredith RM. 2019. Early postnatal development of pyramidal neurons across layers of the mouse medial prefrontal cortex. Sci Rep 9: 5037.

Lomvardas S, Maniatis T. 2016. Histone and DNA Modifications as Regulators of Neuronal Development and Function. Cold Spring Harb Perspect Biol 8: a024208.

Lu L, Liu X, Huang W-K, Giusti-Rodríguez P, Cui J, Zhang S, Xu W, Wen Z, Ma S, Rosen JD, et al. 2020. Robust Hi-C Maps of Enhancer-Promoter Interactions Reveal the Function of Non-coding Genome in Neural Development and Diseases. Molecular Cell 79: 521–534.e15.

Luo L, O’Leary DDM. 2005. Axon Retraction and Degeneration in Development and Disease. Annual Review of Neuroscience 28: 127–156.

Malik AN, Vierbuchen T, Hemberg M, Rubin AA, Ling E, Couch CH, Stroud H, Spiegel I, Farh KK-H, Harmin DA, et al. 2014. Genome-wide identification and characterization of functional neuronal activity–dependent enhancers. Nat Neurosci 17: 1330–1339.

Mall M, Kareta MS, Chanda S, Ahlenius H, Perotti N, Zhou B, Grieder SD, Ge X, Drake S, Euong Ang C, et al. 2017. Myt1l safeguards neuronal identity by actively repressing many non-neuronal fates. Nature 544: 245–249.

Manukyan A, Kowalczyk I, Melhuish TA, Lemiesz A, Wotton D. 2018. Analysis of transcriptional activity by the Myt1 and Myt1l transcription factors. Journal of Cellular Biochemistry 119: 4644–4655.

Moudgil A, Wilkinson MN, Chen X, He J, Cammack AJ, Vasek MJ, Lagunas T, Qi Z, Lalli MA, Guo C, et al. 2020. Self-Reporting Transposons Enable Simultaneous Readout of Gene Expression and Transcription Factor Binding in Single Cells. Cell 182: 992–1008.e21.

Mulvey B, Lagunas T, Dougherty JD. 2021. Massively Parallel Reporter Assays: Defining Functional Psychiatric Genetic Variants Across Biological Contexts. Biological Psychiatry 89: 76–89.

Naruse Y, Aoki T, Kojima T, Mori N. 1999. Neural restrictive silencer factor recruits mSin3 and histone deacetylase complex to repress neuron-specific target genes. Proceedings of the National Academy of Sciences 96: 13691–13696.

Nishibuchi G, Shibata Y, Hayakawa T, Hayakawa N, Ohtani Y, Sinmyozu K, Tagami H, Nakayama J. 2014. Physical and Functional Interactions between the Histone H3K4 Demethylase KDM5A and the Nucleosome Remodeling and Deacetylase (NuRD) Complex *. Journal of Biological Chemistry 289: 28956–28970.

Nitarska J, Smith JG, Sherlock WT, Hillege MMG, Nott A, Barshop WD, Vashisht AA, Wohlschlegel JA, Mitter R, Riccio A. 2016. A Functional Switch of NuRD Chromatin Remodeling Complex Subunits Regulates Mouse Cortical Development. Cell Rep 17: 1683–1698.

Noack F, Vangelisti S, Raffl G, Carido M, Diwakar J, Chong F, Bonev B. 2022. Multimodal profiling of the transcriptional regulatory landscape of the developing mouse cortex identifies Neurog2 as a key epigenome remodeler. Nat Neurosci 25: 154–167.

Oki S, Ohta T, Shioi G, Hatanaka H, Ogasawara O, Okuda Y, Kawaji H, Nakaki R, Sese J, Meno C. 2018. ChIP-Atlas: a data-mining suite powered by full integration of public ChIP-seq data. EMBO reports 19: e46255.

Olson JM, Asakura A, Snider L, Hawkes R, Strand A, Stoeck J, Hallahan A, Pritchard J, Tapscott SJ. 2001. NeuroD2 Is Necessary for Development and Survival of Central Nervous System Neurons. Developmental Biology 234: 174–187.

Pataskar A, Jung J, Smialowski P, Noack F, Calegari F, Straub T, Tiwari VK. 2016. NeuroD1 reprograms chromatin and transcription factor landscapes to induce the neuronal program. The EMBO Journal 35: 24–45.

Romm E, Nielsen JA, Kim JG, Hudson LD. 2005. Myt1 family recruits histone deacetylase to regulate neural transcription. J Neurochem 93: 1444–1453.

Shepherd GM, Rowe TB. 2017. Neocortical Lamination: Insights from Neuron Types and Evolutionary Precursors. Front Neuroanat 11: 100.

Skene PJ, Henikoff S. 2017. An efficient targeted nuclease strategy for high-resolution mapping of DNA binding sites ed. D. Reinberg. eLife 6: e21856.

Stiles J, Jernigan TL. 2010. The Basics of Brain Development. Neuropsychol Rev 20: 327–348.

Subramanian A, Tamayo P, Mootha VK, Mukherjee S, Ebert BL, Gillette MA, Paulovich A, Pomeroy SL, Golub TR, Lander ES, et al. 2005. Gene set enrichment analysis: A knowledge-based approach for interpreting genome-wide expression profiles. PNAS 102: 15545–15550.

Trevino AE, Sinnott-Armstrong N, Andersen J, Yoon S-J, Huber N, Pritchard JK, Chang HY, Greenleaf WJ, Paşca SP. 2020. Chromatin accessibility dynamics in a model of human forebrain development. Science 367: eaay1645.

Tutukova S, Tarabykin V, Hernandez-Miranda LR. 2021. The Role of Neurod Genes in Brain Development, Function, and Disease. Frontiers in Molecular Neuroscience 14. https://www.frontiersin.org/articles/10.3389/fnmol.2021.662774 (Accessed September 7, 2022).

Vierbuchen T, Ostermeier A, Pang ZP, Kokubu Y, Südhof TC, Wernig M. 2010. Direct conversion of fibroblasts to functional neurons by defined factors. Nature 463: 1035–1041.

Wöhr M, Fong WM, Janas JA, Mall M, Thome C, Vangipuram M, Meng L, Südhof TC, Wernig M. 2022. Myt1l haploinsufficiency leads to obesity and multifaceted behavioral alterations in mice. Molecular Autism 13: 19.

Yasumura A, Omori M, Fukuda A, Takahashi J, Yasumura Y, Nakagawa E, Koike T, Yamashita Y, Miyajima T, Koeda T, et al. 2019. Age-related differences in frontal lobe function in children with ADHD. Brain and Development 41: 577–586.

Yousefi S, Deng R, Lanko K, Salsench EM, Nikoncuk A, van der Linde HC, Perenthaler E, van Ham TJ, Mulugeta E, Barakat TS. 2021. Comprehensive multi-omics integration identifies differentially active enhancers during human brain development with clinical relevance. Genome Medicine 13: 162.

Yuan W, Ma S, Brown JR, Kim K, Murek V, Trastulla L, Meissner A, Lodato S, Shetty AS, Levin JZ, et al. 2022. Temporally divergent regulatory mechanisms govern neuronal diversification and maturation in the mouse and marmoset neocortex. Nat Neurosci 25: 1049–1058.

